# Metagenomic selections reveal diverse antiphage defenses in human and environmental microbiomes

**DOI:** 10.1101/2025.02.28.640651

**Authors:** Luis Rodriguez-Rodriguez, James Pfister, Liam Schuck, Arabella E. Martin, Luis M. Mercado-Santiago, Vincent S. Tagliabracci, Kevin J. Forsberg

## Abstract

To prevent phage infection, bacteria have developed an arsenal of antiphage defense systems. Using functional metagenomic selections, we identified new examples of these systems from human fecal, human oral, and grassland soil microbiomes. Our antiphage selections in *Escherichia coli* revealed over 200 putative defenses from 14 diverse bacterial phyla, highlighting the broad phylogenetic interoperability of these systems. Many defense systems were unrecognizable based on sequence or predicted structure, so could only be identified via functional assays. In mechanistic studies, we show that some defense systems encode nucleases that only degrade covalently modified phage DNA, but which accommodate diverse chemical modifications. We also identify outer membrane proteins that prevent phage adsorption and a set of previously unknown defense systems with diverse antiphage profiles and modalities. Most defenses acted against at least two phages, indicating that broadly acting systems are widely distributed among non-model bacteria.

## Introduction

Bacteria must withstand the near constant threat of infection by bacteriophages (or, simply, phages). To block infection, bacteria have devised various antiphage defense systems, which, in turn, have spurred phages to develop counter-defenses to overcome bacterial immunity and re-establish infection^1-4^. This arms race stimulates genetic diversification in both host and virus in nearly all microbial ecosystems on Earth. Thus, many defense and counter-defense systems likely await discovery^2^. Previously described defense and counter-defense systems have become ubiquitous tools in molecular biology, are evolutionary antecedents to some eukaryotic innate immune factors, and often dictate phage infection outcomes^5-8^.

In recent years, over 100 new antiphage defense systems have been identified among genes of unknown function in bacterial genomes^9,10^. Most were found based on genetic proximity to known defenses^2^. Though this ‘guilt-by-association’ search strategy has proven effective, it will miss defense systems in unfamiliar genomic loci. Indeed, machine learning models predict that dozens of new antiphage defenses exist in just *Escherichia coli*, one of the best-studied bacteria^11^. The discrepancy between catalogued and actual defense systems is probably higher for non-model bacteria, as their pangenomes are less characterized and their phages poorly described^12^. Yet, these bacteria are likely targets of many emerging phage technologies^13-15^. To address this discrepancy, we used functional metagenomic selections to identify antiphage defenses from natural bacteria in diverse human-associated and environmental microbiomes.

Functional selections exploit the strong fitness advantages conferred by defense systems to recover them from large libraries of plasmids that each express a different DNA insert^16-18^. Rare DNA inserts that encode a defense system can confer a survival phenotype, so will theoretically outcompete other inserts in a library upon phage challenge. Because functional selections do not use sequence to make predictions, they are well-suited to interrogate genes of unknown function for activities like antiphage defense. Indeed, using only *E. coli* genomes, this approach identified ten new defense systems in loci previously unknown to encode defenses^17^. More recently, a single library selected against a single phage revealed an antiphage system without homology to known defenses^18^. Based on these initial findings, we developed functional selections that could be performed at scale, enabling us to probe diverse ecosystems with multiple phages to reveal many new defense systems.

We surveyed nine metagenomes from human fecal, human oral, and grassland soil microbiomes for antiphage defenses against a panel of seven *E. coli* phages. These 63 selections produced a total of 203 unique DNA inserts with an estimated 88% true discovery rate. The DNA inserts originate from 14 diverse bacterial phyla yet act in *E. coli* to defend against a panel of *E. coli* phages. Nucleases were particularly common in our dataset, suggesting that this form of phage defense may readily traverse phylogenetic barriers. Over 40% of our DNA inserts lacked any recognizable antiphage defense genes, hinting at dozens of potentially novel systems. By interrogating a subset of these inserts, we define nine new antiphage defenses, many that originate in non-model bacteria. Our findings include undescribed nucleases that act against modified phage DNA, β-barrel outer membrane proteins that prevent phage adsorption, and a set of unknown genes with diverse antiphage properties.

### Functional metagenomic selections to uncover antiphage defense systems

We designed functional selections to isolate rare antiphage defenses from complex metagenomic DNA libraries in *E. coli* (Table S1). Unlike previous studies, we used plasmid libraries that express small DNA inserts (roughly 1kb to 6kb), which are long enough to encode complete systems but short enough to enable the rapid identification of causal genes (Table S1)^17-21^. By simplifying the genetics of defense system discovery, we could scale up our selections to survey many microbiomes for defenses against many phages. Our approach employed iterative selections where phage resistance is monitored via colony formation in soft agar overlays (Figure 1A). In this scheme, bulk metagenomic plasmids are purified from colonies that withstand one round of phage selection, transformed into fresh cells, and subjected to a second round of infection. This enriches for phage defense encoded by metagenomic plasmids over resistance that arises due to genomic mutation, enabling the recovery of *bona fide* defense systems.

**Figure 1.**
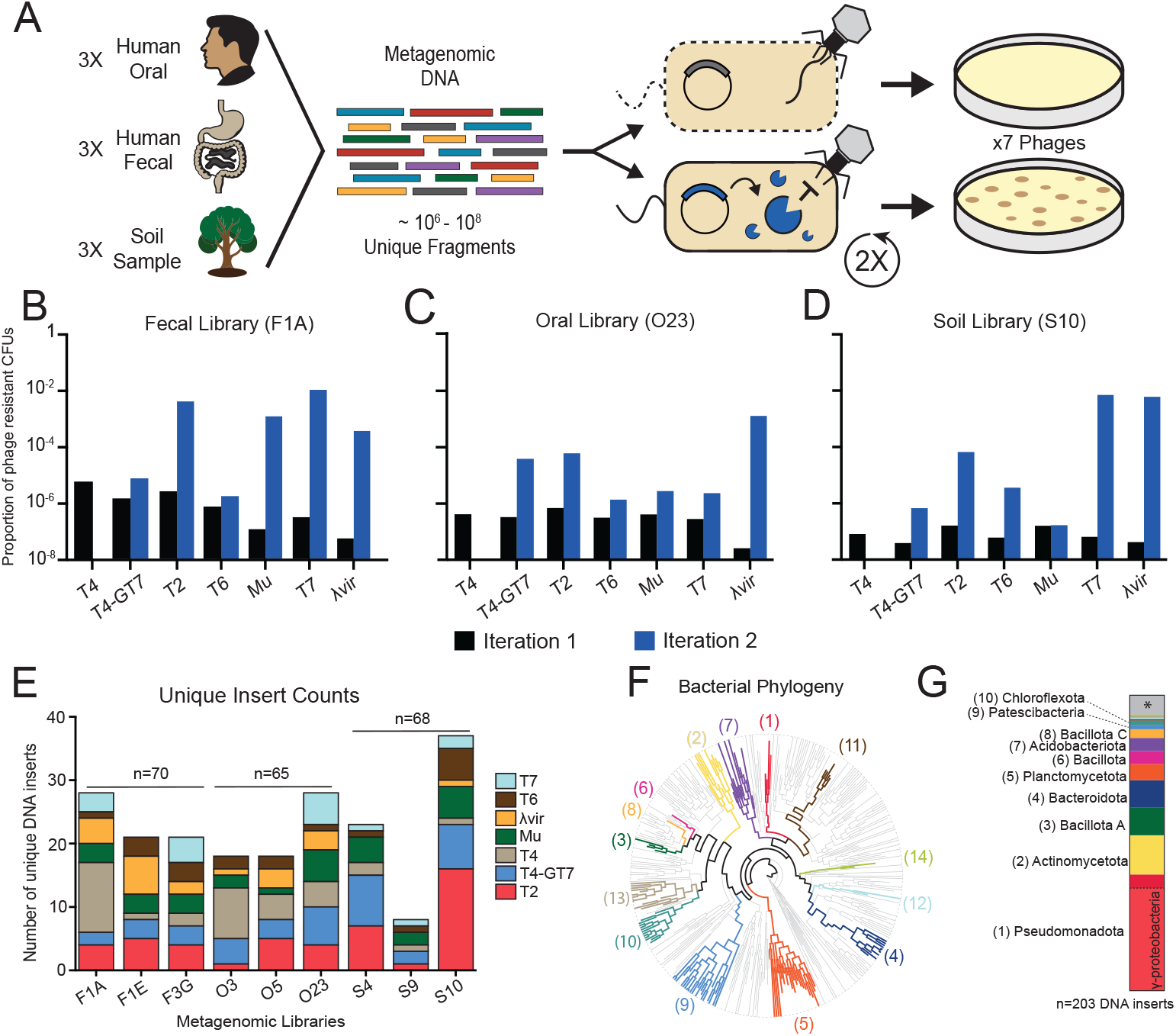
Functional metagenomic selections reveal antiphage defenses. (A) Scheme for functional metagenomic selections. Nine DNA libraries in *E. coli* were subjected to infection with a panel of seven phages. (B-D) Proportion of phage-resistant colony forming units (CFUs) after two rounds of iterative phage selection. Proportions for (B) Fecal (F1A), (C) Oral (O23), and (D) Soil (S10) after the first and second iteration are shown in black and blue, respectively. An elevated proportion of phage resistance suggests a enrichment for phage defense elements. (E) Distribution of unique DNA inserts from 60 independent selections. Each bar represents a different metagenomic library, while the colored layers indicate the phage used for selection. (F) A phylogenetic tree of known bacteria. Source phyla for putative phage defense inserts are depicted in bold and color; each color represents a different phylum. Each tip represents a bacterial class and branch lengths are cropped at a defined radius. Colors and numbers correspond to the stacked bar chart in (G). (G) Predicted phyla-of-origin for all 203 putative phage defense inserts. The ten most prevalent phyla are named and numbered to the left of the stacked bar chart. The 13 defense inserts that lack a confident phylum-level prediction are indicated in gray and with an asterisk. Class-level predictions of γ-proteobacteria are depicted as a subset of their parent phylum, Pseudomonadota, below the dashed line.

To optimize our selection scheme, we used a fecal metagenomic library (named F1A) and two *E. coli* phages: the Tequatrovius phage T4 and a lytic-only mutant of the Lambdavirus, λ_vir_. In the case of T4, phage-resistant colonies emerged from initial selections using F1A but were not seen for selections with an empty vector control (Table S2). When plasmids from F1A colonies were isolated and re-tested for phage defense, 19 of 20 provided strong protection against T4 (95%). For selections with λ_vir_, we observed similar rates of phage resistance for F1A and our empty vector control, regardless of the multiplicity of infection (MOI) used. This suggests that many λ_vir_-resistant colonies did not arise due to the action of metagenomic DNA. Indeed, only one of 16 plasmids conferred λ_vir_ resistance when purified and re-tested in a fresh host background (Figure S1, Table S2). Thus, one round of phage infection was sufficient to reveal metagenomically encoded defenses against phage T4, but not against λ_vir_. However, a second round of selection with λ_vir_ enriched for plasmid-encoded defenses well above background (Figure S1, table S2). We therefore used iterative selections to survey metagenomes for their antiphage defenses.

To identify phage defense elements, we surveyed three metagenomic libraries from grassland soils^19^, three from oral swabs of Yanomami Amerindians^20^, and three from fecal samples of peri-urban residents of Lima, Peru^21^ (Table S1). These libraries were estimated to contain between 1.3×10^6^ and 9.0×10^6^ unique plasmids, each with a different metagenomic DNA insert^19-21^. We subjected each library to iterative selection for defense against one of seven well-studied phages using soft agar overlays (Figure 1A, Table S3). In the case of phage T4, only one round of infection was required because empty vector selections were devoid of phage resistance (Table S4). For all other phages, two rounds of selection were needed, as initial infections revealed similar rates of resistance for libraries and empty vector controls (Table S4). For most selections, the second round of selection substantially increased rates of phage resistance compared to the first. This indicates that phage defense was mediated by our plasmid libraries (Figures 1B-1D, Tables S4, S5). From these defense-enriched selections, we purified plasmids, performed long-read sequencing, and assembled metagenomic DNA inserts that putatively drive our antiphage defense phenotypes.

### Phage defense systems are encoded by diverse metagenomes

We assembled high confidence DNA sequences from 60 of our 63 independent selections, yielding 203 putative phage defense inserts from soil, fecal, and oral metagenomic libraries (Figure 1E, Table S6). In aggregate, each biome produced similar numbers of unique inserts and included putative defense systems against each phage surveyed (Figure 1E). Taxonomic predictions revealed that 72 defense inserts (35%) came from members of the γ-proteobacteria, a bacterial class that includes our selection host, *E. coli* (Figures 1F, 1G, and Table S6). Though γ-proteobacteria were the most common bacterial class in our phage defense dataset, they represent only 2.6% of bacteria in our soil, fecal and oral metagenomes, based on a reanalysis of 16S rDNA datasets from these exact samples (Table S7)^19-22^. This overrepresentation indicates that antiphage defenses from γ-proteobacteria are more likely to function in *E. coli* than defense systems from other, more diverged bacteria. This may be due to some defense systems that only retain function across short phylogenetic distances^23^.

Surprisingly, most DNA inserts came from bacterial phyla distinct from *E. coli* (110/203, or 54%, arose outside the Pseudomonadota). In total, DNA from 14 different bacterial phyla prevented phage infection in *E. coli*, including examples from gram-positive, gram-negative, and candidate phyla radiation bacteria (Figures 1F, 1G, and Table S6). To assess potential phylogenetic bias among these inserts, we compared our phage defense and 16S datasets, omitting γ-proteobacteria from the analysis. We observed a significant correlation between each phylum’s abundance in our phage defense dataset and its abundance as measured by 16S rDNA sequencing (Figure S2). Thus, once close relatives of *E. coli* are ignored, phylogenetic relationships do not substantially bias whether our defense inserts can operate in this bacterial host. This broad interoperability suggests that many antiphage defenses target conserved or recurrent features of phage infection. This may explain why some defense genes move frequently between distantly related bacteria^24^.

Known defense systems tend to cluster in discrete genomic loci called defense islands^25^. Thus, we looked for signatures of defense islands in our dataset. Over half of our phage defense inserts had strong nucleotide identity to at least one sequence in NCBI’s nt database, allowing us to examine their genomic context. In almost all these queries, the insert mapped within 10kb of a known defense gene, indicating a defense island origin (Table S8). This suggests that our dataset is enriched for antiphage defenses. Indeed, most DNA inserts (121/203, 60%) encoded at least one open reading frame (ORF) with homology to a known defense protein (e-value < 10^−5^). In many cases, these inserts did not encode complete defense systems but rather contained homologs of isolated defense genes or candidate systems (Supplemental Dataset 1)^9,10,26-28^. We annotated putative nuclease domains in over half of DNA inserts (55%), indicating a strong enrichment for nucleic acid degradation in our dataset (Table S6). We also identified 82 sequences without a defense-associated gene (40% of inserts). In summary, our dataset exhibits the expected hallmarks of defense-associated sequences and likely contains novel antiphage defense systems.

To evaluate the robustness of our phage defense dataset, we individually tested 73 diverse DNA inserts for defense against the phage on which they were originally identified (Table S6). When expressed from a plasmid in *E. coli*, we observed reproducible phage defense for 64 of 73 inserts (88%), indicating that most DNA inserts in our dataset confer *bona fide* antiphage defense (Figures 2A, S3). We then re-tested these 64 sequences for defense against the full seven-phage panel, revealing 53 sequences that conferred >100-fold protection against at least two phages (Figure 2A). The majority of these multi-phage defenses provided protection against phages with different receptors and from different families (Table S3). Thus, we find many defense systems in commensal and environmental bacteria that confer potent and broad-spectrum protection against *E. coli* phages.

**Figure 2.**
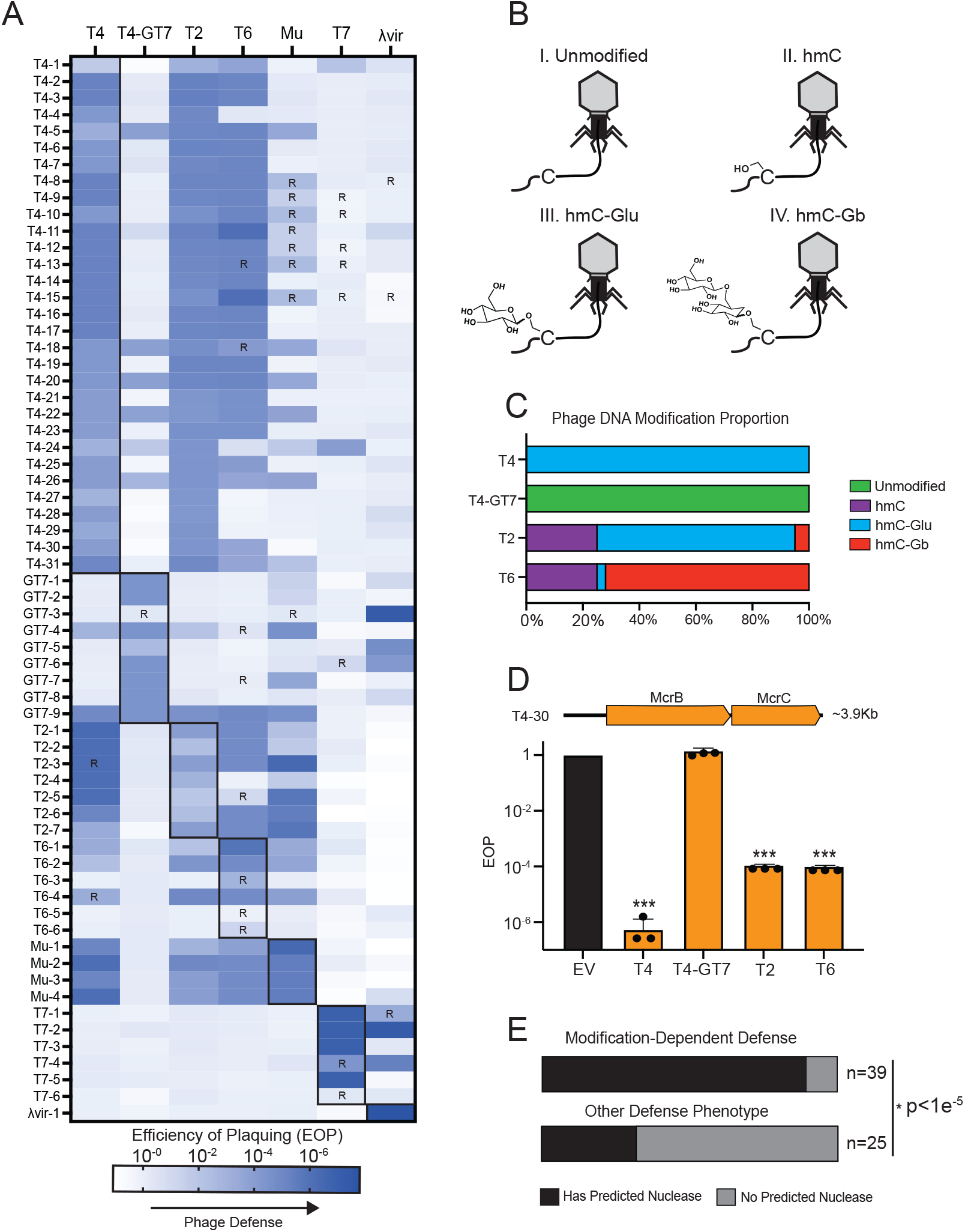
Metagenomic inserts confer broad-spectrum phage defense. (A) Heat map showing efficiency of plaquing for a set of 64 validated DNA inserts against a panel of seven phages. Bold outlines depict infections with the phage used in an insert’s initial discovery. R; reduced plaque size. (B) Cartoon of four T-even phages, each with a different covalent modification on cytosines. I. Unmodified cytosine, II. Hydroxymethyl cytosine (hmC), III. Glucosyl hydroxymethyl cytosine (hmC-Glu) and, IV. Gentiobiose hydroxymethyl cytosine (hmC-Gb). (C) Stacked bar chart depicting the proportion of cytosine modifications for each of the T-even phages, per ref. 29. (D) Efficiency of plaquing (EOP) for T-even phages against insert T4-30, which encodes a homolog of the type IV restriction system McrBC, but no other ORFs; *** denotes p≤0.0001 vs T4-GT7, one way ANOVA, n=3. EV, empty vector; error bars depict standard deviation. (E) Stacked bar chart analysis showing the number of nuclease-encoding inserts associated with modification dependent defense phenotypes versus others; p<1e^-5^, Fisher Exact Test.

### Modification-Dependent Nucleases Block Phage Infection

In some cases, the phenotypic profile of a phage defense insert enabled us to predict a putative defense mechanism. For instance, many defense inserts from our metagenomic selections provide protection against the phages T2, T4, and T6, but not the T4-derived mutant T4-GT7 (Figure 2A). T-even phages covalently modify cytosines to protect their DNA from bacterial nucleases, whereas T4-GT7 is incapable of this modification^29-31^ (Figures 2B, 2C). Thus, we hypothesize that these defense inserts recognize modified DNA as a ‘non-self’ marker of phage infection. Indeed, we find multiple homologs of the type IV restriction system McrBC, which uses a modification-dependent nuclease to target phage genomic DNA^32^. As expected, an insert encoding an McrBC-like system blocked infection by T2, T4, and T6, but not T4-GT7 (Figure 2D). Further supporting a DNA-centric immune strategy, we find at least one putative nuclease on 35 of the 39 inserts that defend against T4 but not T4-GT7, a significant enrichment compared to the other inserts in our dataset (p<1e^-5^, Fisher’s Exact Test, Figure 2E, Table S6). Like the recently described antiphage effector BRIG1, these enzymes may include previously unknown defenses that act against phages with modified DNA nucleotides^18^.

During our work, two nucleases, HEC-06 and PD-T4-3, were reported to block T4 infection when expressed heterologously in *E. coli*^17,28^. We found homologs of each nuclease among our inserts and selected two for further study (named HEC-06_Bact_ and PD-T4-3_Capno_ to reflect their predicted host genera, *Bacteroides* and *Capnocytophaga*). HEC-06_Bact_ and PD-T4-3_Capno_ share 22% and 48% amino acid identity with their cognate nucleases, respectively. To test whether these nucleases have a role in phage defense, we cloned each behind an inducible promoter, expressed them in *E. coli*, and challenged each strain with a series of T-even phages. Compared to GFP controls, both predicted nucleases provided strong defense against at least two T-even phages, with PD-T4-3_Capno_ acting at very low expression levels (i.e. in the absence of inducer, Figures 3A, 3B). Regardless of expression level, neither gene could protect against phage T4-GT7, which lacks cytosine modification^30,31^ (Figures 3A, 3B). This suggests that both genes recognize modified phage DNA to elicit phage defense. To test this hypothesis further, we purified PD-T4-3_Capno_ and monitored its ability to degrade different phage genomic DNA substrates. The enzyme readily degraded modified T4 genomic DNA *in vitro* but showed no activity against unmodified DNA purified from T4-GT7 (Figure 3C). These results suggest that DNA modifications may activate PD-T4-3 family nucleases, which could explain their ability to block phage infection.

**Figure 3.**
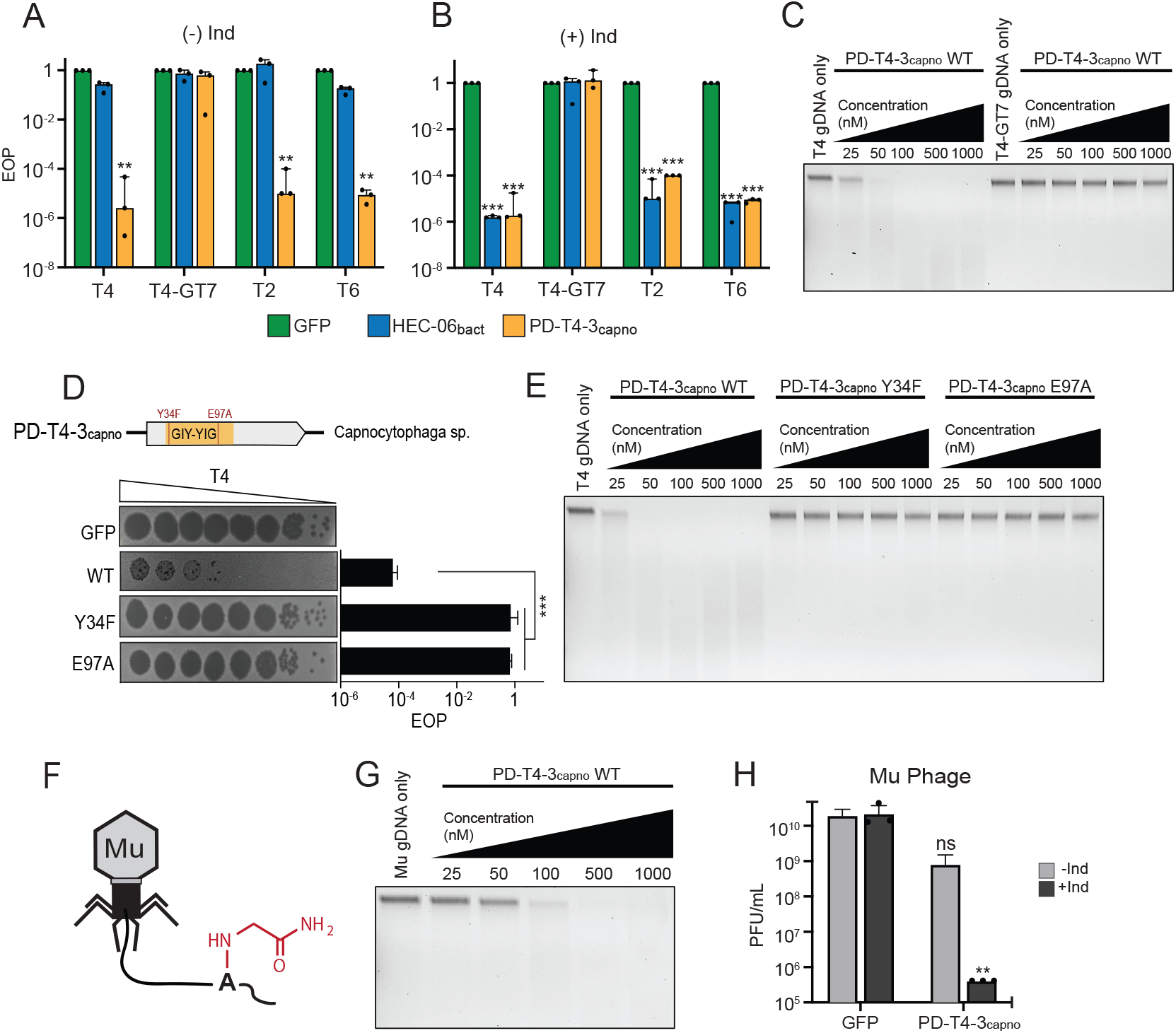
Nucleases cleave modified phage genomes to elicit defense. (A-B) Efficiency of plaquing for HEC-06_Bact_ and PD-T4-3_Capno_ against T-even phages compared to GFP controls; (A) no inducer, (B) with inducer; significance: ** p≤0.001, *** p≤0.0001 (one way ANOVA vs T4-GT7). (C) Purified PD-T4-3_Capno_ incubated with T4 and T4-GT7 genomic DNA for 10’ at different concentrations; DNA was stained with ethidium bromide and visualized under UV light after agarose gel electrophoresis. (D) Predicted catalytic residues for PD-T4-3_Capno_ and their impact on infection via plaque assay; *** p≤0.0001 (one way ANOVA vs WT). (E) The degradation of T4 genomic DNA *in vitro* by PD-T4-3_Capno_ and its mutants. Reaction products were analyzed as in (C). (F) Cartoon of phage Mu showing the addition of a carbamoyl methyl group to the N6 position of adenine. (G) In vitro degradation of Mu genomic DNA by purified PDT4-3_Capno_ at different concentrations. Reaction products were analyzed as in (C). (H) Defense of PD-T4-3_Capno_ against phage Mu compared to a GFP control; PFU = plaque forming units, (+/-) Ind indicates whether expression was induced; ** p≤0.001, ns = no significance (Welch’s T-test vs condition-match GFP control). All experiments in biological triplicate; error bars depict standard deviation.

PD-T4-3 was the most common annotation in our dataset (Table S9) and these homologs were almost always associated with modification-dependent phage defense (defined as restriction of T4 but not T4-GT7, Table S6). To test if nuclease activity was required for defense, we mutated conserved residues in the predicted active site of PD-T4-3_Capno_. These mutants failed to defend against T4 *in vivo* (Figure 3D) and could not degrade T4 genomic DNA *in vitro* (Figure 3E). Despite acting against T4, PD-T4-3_Capno_ was originally recovered from selections against phage Mu. Mu does not modify cytosines but instead adds a carbamoyl methyl group to the N6 position of adenine^33,34^ (Figure 3F). Based on this finding, we tested whether PD-T4-3_Capno_ could degrade genomic DNA from phage Mu. We observed efficient degradation at high protein concentrations (0.1 to 1 µM) but almost no activity at lower levels (Figure 3G). A similar trend was seen *in vivo*, as a leaky expression of PD-T4-3_Capno_ provided only moderate defense against phage Mu (unlike the strong defense against phage T4, Figure 3A), but substantial defense was observed upon PD-T4-3_Capno_ induction (Figure 3H). These results suggest that PD-T4-3_Capno_ exhibits some plasticity to recognize different nucleotide modifications and elicit phage defense, perhaps explaining its widespread prevalence across unrelated bacterial genomes^17^.

### Beta-Barrel Outer Membrane Proteins Block Phage Adsorption

We noticed that several DNA inserts from selections against phage T7 also provided defense against λ_vir_ (Figure 2A, lower right). We were intrigued by this shared defense phenotype, especially because these phages belong to different families, recognize different primary receptors, and employ different replication strategies^35,36^. To identify potential defense genes, we removed the start codon from each ORF on the DNA insert T7-2, which exhibited the strongest T7 and λ_vir_ defense. We then challenged these mutants with T7. This experiment revealed an 8-stranded, β-barrel outer membrane protein (OMP) that was necessary for phage defense (Figure 4A), which we named OMP_Tell-2_ to reflect its predicted host genus, *Telluria*, and its parent insert, T7-2. Two other DNA inserts conferred dual defense against T7 and λ_vir_, T7-1 and T7-4 (Figure 2A). These inserts encoded 8-stranded and 12-stranded β-barrel OMPs, which we named OMP_Tell-1_ and OMP_Acin-4_, respectively. When each OMP was expressed in isolation, they provided more than 10^4^-fold protection against T7 infection and up to 6×10^5^-fold protection against λ_vir_ (Figure 4B). Thus, these three OMPs are each sufficient for phage defense.

**Figure 4.**
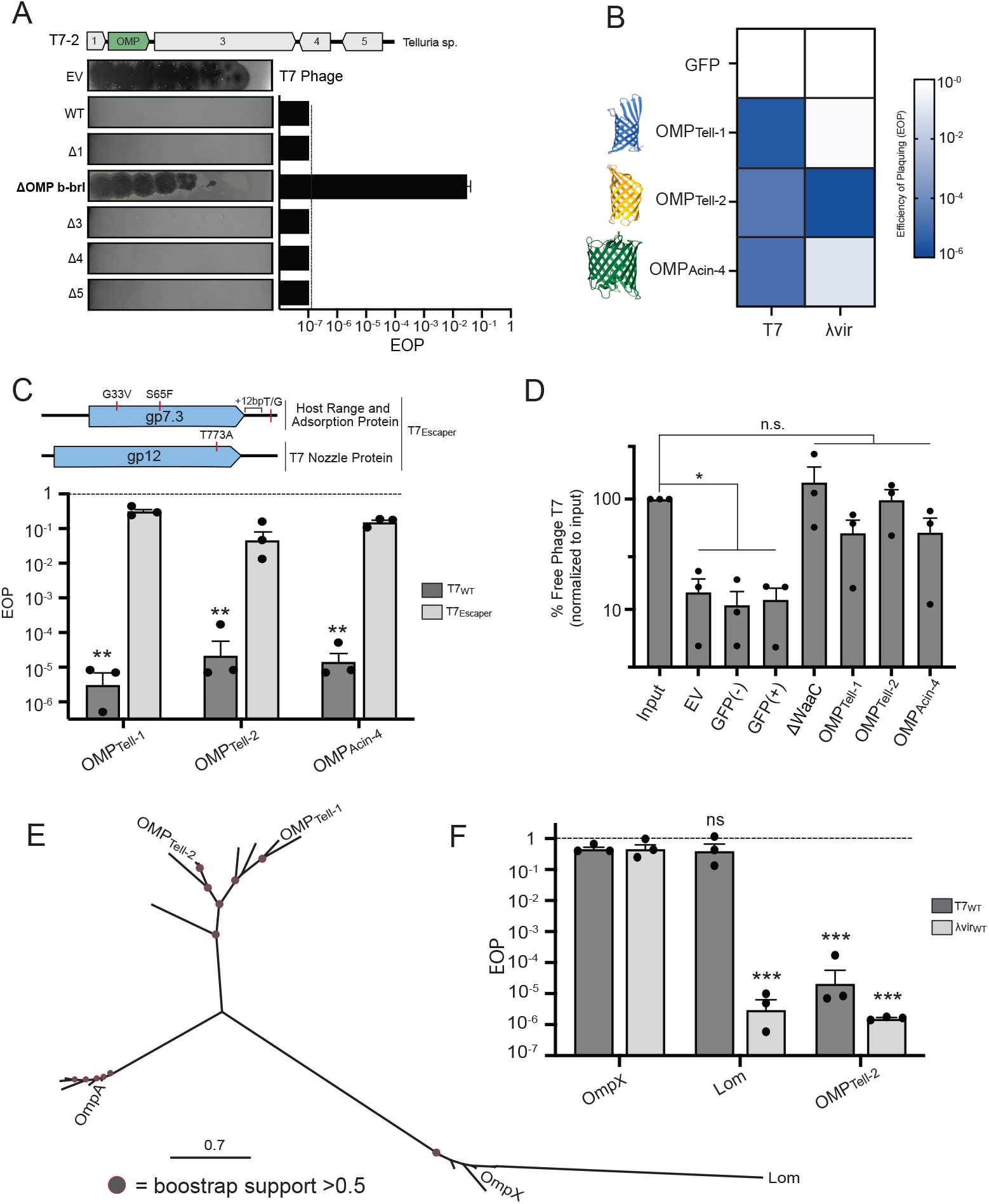
Outer membrane proteins block phage adsorption. (A) A gene encoding a predicted OMP is necessary for phage defense by the T7-2 insert. A Δ symbol indicates that the start codon of indicated gene was deleted. The bar chart indicates the EOP for each genotype upon T7 challenge. Solid line indicates limit of detection. (B) Heat map depicting EOP for GFP, OMP_Tell-1_, OMP_Tell-2_, and OMP_Acin-4_ when challenged with T7 and λvir. Structures of each OMP predicted by Alphafold3 are depicted without their signal peptide. (C) Top, missense mutants in T7_Escaper_ mapped to two loci, encoding genes associated with phage adsorption and DNA injection. Bottom, EOP for T7_WT_ and T7_Escaper_ against OMP_Tell-1_, OMP_Tell-2_, and OMP_Acin-4_; ** p≤0.001 (Welch’s T-test vs T7_WT_). (D) T7-phage adsorption assay. The y-axis depicts free, unadsorbed phage following incubation with *E. coli* cells of the indicated genotypes. Asterisks denote p<0.05, T-test with Bonferroni-Holm correction for multiple hypotheses. (E) An unrooted phylogenetic tree depicting various OMPs and their homologs. Red outlines on gray circles depict nodes with bootstrap support > 0.5. (F) EOP for OmpX, Lom, and OMP_Tell-2_ against T7_WT_ and λ_vir_; *** p≤0.0001, ns = no significance (one way ANOVA vs condition-matched OmpX control). Dashed lines in (C) and (F) indicate infectivity against a GFP control, defining and EOP value of 1. All experiments performed in biological triplicate; error bars depict standard deviation.

During these experiments, we observed rare T7 plaques that formed during infections of OMP_Acin-4_ expressing cells. We isolated, propagated, and sequenced one of these plaques to identify potential mutations that enabled T7 infection. This phage, T7_Escaper_, contained multiple missense mutations across the genes *gp7*.*3* and *gp12*, which encode T7’s host adsorption and nozzle proteins, respectively. These proteins physically interact to promote adsorption and DNA injection during T7 infection^37-39^. Interestingly, this T7_Escaper_ phage could infect *E. coli* expressing OMP_Tell-1_, OMP_Tell-2_, and OMP_Acin-4_ similarly well, despite being recovered from only from the latter strain (Figure 4C). This suggests that all three OMPs employed a similar mechanism of T7 defense, which we suspected may be related to phage adsorption. To test this idea, we measured the proportion of T7 phages that adsorb to *E. coli* expressing each OMP, empty vector and GFP controls, and a lipopolysaccharide biosynthesis mutant (ΔWaaC) known to impair T7 adsorption^36^. Following centrifugation, most phages were removed from the supernatant of empty vector and GFP controls, indicating a co-sedimentation with these cells and thus strong adsorption of T7 to these strains (Figure 4D). In contrast, the proportion of free phages was statistically indistinguishable from input for the ΔWaaC mutant and each OMP-expressing strain (Figure 4D). This suggests that the expression of these OMPs prevents T7 infection by blocking phage adsorption, as occurs for the ΔWaaC mutant^36^.

We next searched for potential homologs of each phage-resistant OMP. Sequence comparisons by BLAST and structural predictions using AlphaFold3 revealed that OMP_Acin-4_ was related to a family of *Acinetobacter* porins with poorly defined activity (Figure S4). Similarly, OMP_Tell-1_ and OMP_Tell-2_ are sequence and structural homologs of the eight-stranded, β-barrel proteins OmpA and OmpX from *E. coli* (Figures 4E, S4). OMP_Tell-1_ and OMP_Tell-2_ are homologous to one another and cluster separately from OmpA and OmpX-like proteins on a phylogenetic tree. We could not establish a phage defense phenotype for OmpA or OmpX but observed strong λ_vir_ defense upon expression of a long-branching OmpX-family protein, named Lom^40^ (<u>L</u>ambda <u>o</u>uter <u>m</u>embrane, Figure 4F). Lom is a β-barrel protein encoded by phage λ that has been associated with adhesion and virulence in some *E. coli* strains but has no other known function^41^. Lom’s strong antiphage defense phenotype in our experiments suggests a potential antiphage role in nature. Consistent with this possibility, Lom is one of the few proteins expressed constitutively from the λ prophage, a characteristic typical of prophage-encoded antiphage defense and superinfection exclusion genes^42-44^.

### Novel Defense Systems

Many confirmed defense inserts lacked ORFs with detectable homology to known defense genes (Figures 2A, S3). To determine if these inserts encoded new phage defenses, we selected six examples for genetic interrogation. We first mutated the start codon of each full-length ORF to reveal genes that impacted our phage defense phenotype (Figure S5). We then tested whether these genes were sufficient for defense in solid and liquid media and against our full panel of phages, revealing four single-gene defenses and two instances of more complex systems (Figures 5A, 5B, S6, S7). We named individual defense genes using standard genetic nomenclature and assigned the names San Juan and Dallas to the more complex defense systems. We then examined each defense gene for its prevalence among bacteria in NCBI, revealing some genes with narrow phylogenetic distributions and others in many bacterial clades (Figure 5C). The defense genes *ldgA* and *ddgA* appear almost exclusively in the Patescibacteria, a massive clade of obligate epibionts previously called the candidate phyla radiation^45^. These genes, along with the Planctomycete-derived *tdgA*, do not have significant sequence or structural homology to proteins of known function, obscuring mechanistic predictions. However, based on their shared phage defense profile, we speculate that *tdgA* and Dallas recognize common features of phage infection, which are distinct from the λ_vir_-specific defense mechanism of the gene *ldgA* (Figure 5B).

**Figure 5.**
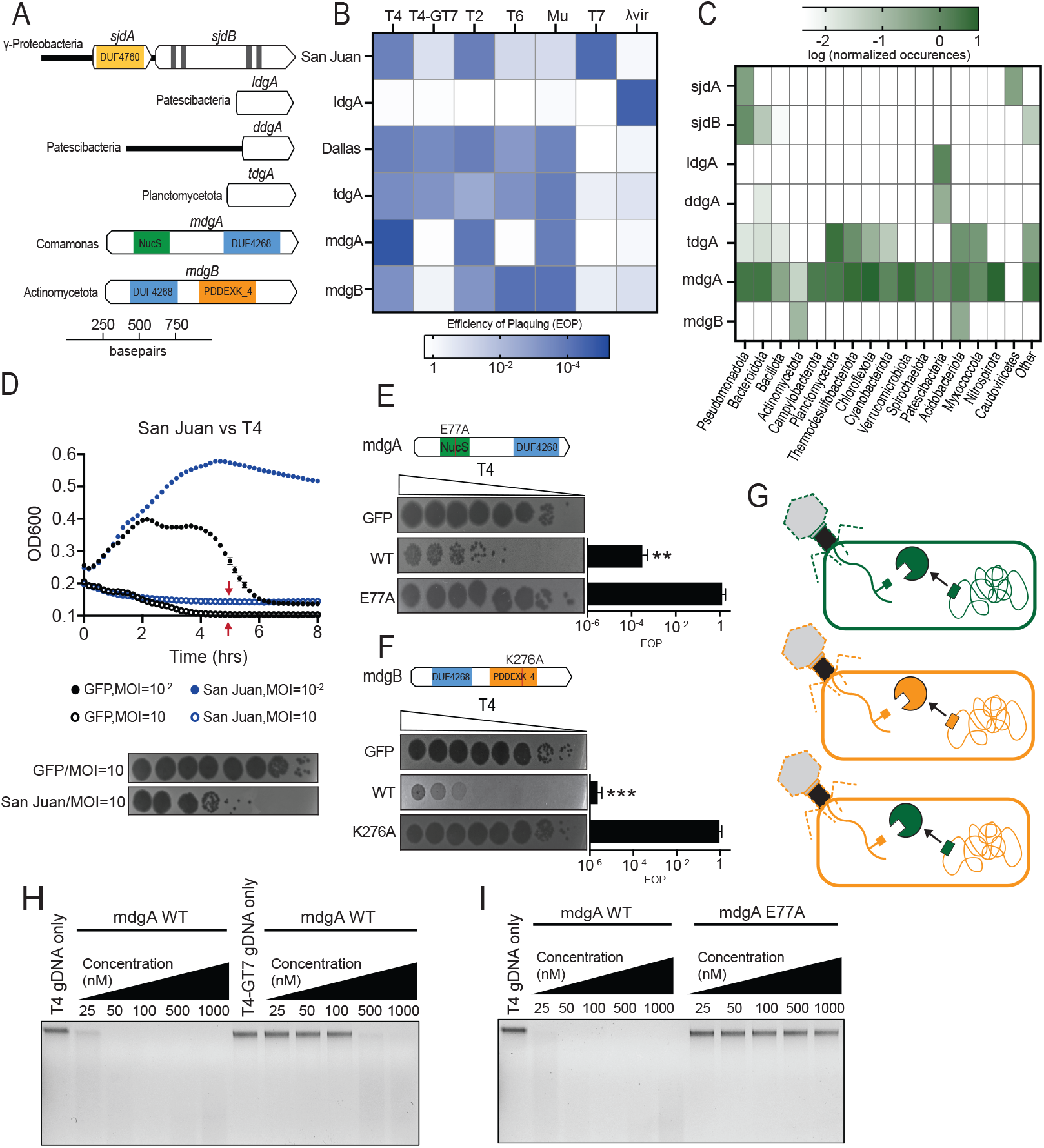
Novel defenses exhibit diverse antiphage profiles and modalities. (A) Cartoon representation of two novel defense systems (San Juan and Dallas) and four novel genes (*IdgA, tdgA, mdgA* and *mdgB*). Acronyms: *sjd*, San Juan Defense; *ldg*, Lambda Defense Gene; *ddg*, Dallas Defense Gene; *tdg*, T-even Defense Gene; *mdg*, Modification Defense Gene. (B) Heat map depicting EOP values relative to GFP controls for each defense against a panel of seven phages. (C) A heat map illustrating the prevalence of defense genes across different phyla. (D) T4 infection of San Juan and GFP expressing *E. coli* cells using MOIs of 0.01 and 10. At the timepoint indicated by red arrows, parallel infections were subsampled and PFUs enumerated by plaque assay (bottom). (E) A predicted catalytic residue in MdgA and its impact on T4 infection as measured via plaque assay; ** p≤0.001 (Welch’s T-test, WT vs mutant). (F) A predicted catalytic residue in MdgB and its impact on T4 infection as measured via plaque assay; *** p≤0.0001 (Welch’s T-test, WT vs mutant). (G) A cartoon model depicting the interoperable defense activity of modification dependent nucleases across different host backgrounds. Green, orange depict lineage-specific phage-bacterial arms races where nucleases recognize and cleave modifications on their target phage genome. This activity is cross-compatible across hosts. (H) Purified MdgA incubated with T4 and T4-GT7 genomic DNA for 10’ at different concentrations. Reaction products were analyzed as in Figure 3C. (I) The degradation of T4 genomic DNA by MdgA and its catalytic mutant. Reaction products were analyzed as in Figure 3C. All experiments performed in biological triplicate; error bars depict standard deviation.

San Juan was recovered from selections against T4 and contains two genes (Figures 5A, S5). Though San Juan exhibits stronger restriction against phage T4 than T4-GT7, we could not identify a nuclease domain within either *sjdA* or *sjdB* (*sjd* = <u>S</u>an <u>J</u>uan <u>d</u>efense). Thus, we do not favor modification-dependent nuclease activity as San Juan’s defense mechanism, like we observe for PD-T4-3_Capno_ (Figure 3). It is unlikely that modified nucleotides play any role in San Juan’s defense activity, as it strongly blocks infection by phage T7, which naturally lacks DNA modifications^36^ (Figure 5B). Instead, four putative transmembrane domains in *sjdB* led us to suspect abortive infection as a potential defense strategy. Abortive infection is a common theme in antiviral defense, referring to the programmed death or dormancy of an individual cell to protect the surrounding population. This results in a density-dependent defense phenotype wherein cells succumb to high-titer infections but withstand low-titer ones. To determine if San Juan acts via abortive infection, we subjected *E. coli* expressing either San Juan or GFP to liquid infection with high and low amounts of phage T4. Consistent with abortive infection, San Juan blocked T4 infection at low MOIs, unlike GFP, but both populations declined rapidly at high MOIs (Figure 5D). We sampled these high-titer populations and measured phage numbers to determine if each cell background supported phage replication. We observed 10^3^-fold higher phage titers in the GFP cells compared to the San Juan background (Figure 5D, bottom). This suggests that the population decline observed for San Juan prevents phage replication, unlike the phage-induced lysis experienced by the GFP population. Thus, San Juan elicits phage defense via canonical abortive infection.

Our selections also revealed new phage defense nucleases. We identified two genes that conferred defense against phage T4, but not T4-GT7, and encoded predicted nuclease domains (Figures 5A, 5B). We tentatively named these genes *mdgA* and *mdgB* to reflect their putative modification-dependent nuclease activity (*mdg* = <u>m</u>odification <u>d</u>efense <u>g</u>ene). Homologs of MdgA were prevalent across many bacterial phyla whereas MdgB was restricted to the common soil phyla Actinomycetota and Acidobacteriota (Figure 5C). Both MdgA and MdgB conferred strong phage defense when expressed in *E. coli*, which was lost when putative catalytic residues in the active sites of each nuclease were swapped for alanine (Figures 5E, 5F). To test for modification-dependent nuclease activity, we purified both wild-type MdgA and its catalytic mutant, MdgA_E77A_. Wild-type MdgA potently degraded T4 genomic DNA but showed significantly reduced activity against DNA from T4-GT7 (Figure 5H). MdgA_E77A_ could not degrade T4 genomic DNA at any tested concentration, indicating that this residue is required for nuclease activity, explaining its importance for phage defense (Figures 5E, 5I). Based on these data, we propose that MdgA and MdgB are modification-dependent antiphage nucleases like PD-T4-3_Capno_ and HEC-06_Bact_ (Figure 3)^17,28^. Together, our data indicate that this class of phage defense genes is highly interoperable. Heterologous nucleases from multiple phyla provide phage defense to *E. coli* in a modification-dependent matter (Figures 2A, 2E, 3G). This suggests that DNA modification is a recurrent phage adaptation that can be recognized by promiscuous antiphage nucleases in many bacterial lineages (Figure 5G).

## Discussion

Metagenomes contain genomic DNA from diverse bacteria, most of which encode phage defense systems. Via high throughput functional selections, we establish that defense systems from 9 metagenomes and 14 phyla can work in *E. coli* to prevent phage infection. The abundance of these phyla in our dataset correlates with their abundance in the original metagenomic samples. Thus, many phage defenses retain function across huge evolutionary distances, regardless of source genome or habitat. Modification dependent nucleases are the most common defense function in our dataset, implying that these enzymes traverse phylogenetic barriers particularly well. This could be due to phages that recurrently select for these nucleases across bacterial lineages, promoting interoperability.

We also find defense inserts without nucleases that function in *E. coli*, including 64 examples from bacteria outside its phylum (Table S6). These include 14 inserts confirmed for phage defense and several where we identify causal genes and systems (Figures 3A, 4B, 5A). Some of these defense genes, like *tdgA*, act against many phages and have homologs in diverse phyla. Thus, even though we understand little about *tdgA*’s defense mechanism, we can surmise that it relies on some conserved or recurrent feature of phage infection. Other genes, like *ldgA* and *ddgA* of the Patescibacteria, retain function across tremendous phylogenetic distance but lack homologs in any other bacterial clade, perhaps due to low rates of inter-phyla gene flow in these bacteria^45,46^. Their discovery from oral and soil metagenomes is surprising, as only one Patescibacteria phage has been described and very few defense systems have been identified in these genomes^47,48^. The only known Patescibacteria phage encodes many T4-like structural proteins, which could help explain why the Dallas defense system protects against T-even phages in our panel.

Three DNA inserts confer protection against the phages T7 and λ_vir_ and each encode an OMP that prevents phage adsorption. These OMPs, OMP_Tell-1_, OMP_Tell-2_, and OMP_Acin-4_, have their closest homologs in regions of bacterial genomes without obvious signatures of phage origin. Thus, they may represent dedicated antiphage defense genes in their natural hosts. Phage defense could also be a moonlighting function, as OMPs can exhibit highly pleotropic activities^49^. Unlike its OMP_Tell_ homologs, Lom is encoded by phage λ and protects only against λ_vir_ among tested phages. Thus, Lom’s activity may have evolved to benefit phages rather than host cells, for instance, via superinfection exclusion. Superinfection exclusion can benefit both lysogens and circulating phages by preventing the latter from adsorbing non-productively to an already infected host^43,44^. Though Lom is expressed from integrated prophages, it does not reach sufficient levels to prevent λ_vir_ from infecting a lysogen in laboratory conditions^42,50^. However, non-productive adsorption can also be detrimental during times of coordinated lysis, when early bursting phages frequently encounter hosts with replicating virus^51^. Lom’s expression increases dramatically during late infection, implying it may act at this stage of infection to prevent non-productive adsorption^42^. Consistent with this kin selection model, Lom provided no defense against the heterotypic phage T7 in our experiments (Figure 4F).

Nucleases were the most common function within our phage defense dataset, with phenotypic profiles suggesting that many of them act against phages with modified DNA genomes. We corroborated this activity for four proteins *in vivo* (HEC-06_Bact_, PD-T4-3_Capno_, MdgA, and MdgB) and demonstrated function *in vitro* for PD-T4-3_Capno_ and MdgA. PD-T4-3_Capno_ is a modification-dependent nuclease with high plasticity, capable of degrading both T4 DNA glucose-modified cytosines and Mu DNA with carbamoylmethyl-modified adenines. This plasticity may explain its widespread distribution, as it could enable defense against various phages with different genomic modifications^17^. MdgA is similarly widespread in bacterial genomes and capable of defending against both T4 and Mu *in vivo*. Though analogous to PD-T4-3_Capno_ in this regard, the proteins share no conservation in nuclease or accessory domains, suggestive of convergent evolution. The presence of DUF4268 domains in both MdgA and MdgB represent a potential substrate recognition motif, especially as this domain co-associates with type IV restriction enzymes across bacterial genomes^52^.

From 60 phage selections in *E. coli*, we recovered 203 metagenomic DNA inserts. Many encoded defense systems that were unrecognizable based on their sequence or predicted structure, so could only be identified via functional assays. Frequently, these defenses were widely distributed and broadly interoperable. This implies that the reservoir of defense genes available to any bacterial species is larger than what circulates within its cataloged pangenome. In nature, phages keep pace with this diversity by continually updating their inventory of anti-defense genes, enabling them to re-establish infection in the face of new barriers^3,4^. Analogously, any phage-based technology for eliminating or modifying natural bacteria will need to embrace diversity to sustain success^53^.

## Acknowledgments

We thank Krzysztof Pawlowski for help performing structural homology predictions and John Schoggins, Julie Pfeiffer, and Alex Meeske for comments on manuscript drafts.

## Author contributions

Conceptualization: KJF; Methodology: LRR, JP, VST, KJF; Investigation: LRR, JP, LS, AEM, LMMS, KJF; Funding acquisition: VST, KJF; Project administration: VST, KJF; Supervision: VST, KJF; Writing – original draft: LRR, KJF; Writing – review & editing: LRR, JP, VST, KJF.

## Data and materials availability

Sequence data associated with this project have been deposited to NCBI under BioProject (accession pending). Code developed as part of this project to investigate or process sequence data are deposited on Zenodo (link pending). All other data are available in the main text or the supplementary materials.

## Supplementary Materials

### Supplementary Methods

**Figure S1.** A second round of phage selection enriches for plasmid-encoded defenses.

**Figure S2.** A phylum’s abundance in the phage defense dataset correlates with its abundance as measured by sequencing 16S rDNA amplicons.

**Figure S3.** Representative plaque assays for 64 validated DNA inserts.

**Figure S4.** Structural predictions by AlphaFold3 for OMP_Tell-1_, OMP_Tell-2,_ and OMP_Acin-4_ aligned with their respective structural homologs.

**Figure S5:** Gene(s) necessary for phage defense in unknown systems.

**Figure S6.** Liquid phage infections with various defenses and controls.

**Figure S7**. Phage defense sufficiency experiments via plaque assays.

**Figure S8**. Phage defense inserts are encoded by diverse metagenomes.

**Supplementary Tables S1 to S11**.

## References

1 Forsberg, K. J. & Malik, H. S. Microbial Genomics: The Expanding Universe of Bacterial Defense Systems. Curr Biol 28, R361–R364 (2018). 10.1016/j.cub.2018.02.053

2 Georjon, H. & Bernheim, A. The highly diverse antiphage defence systems of bacteria. Nature Reviews Microbiology 21, 686–700 (2023). 10.1038/s41579-023-00934-x

3 Mayo-Muñoz, D., Pinilla-Redondo, R., Camara-Wilpert, S., Birkholz, N. & Fineran, P. C. Inhibitors of bacterial immune systems: discovery, mechanisms and applications. Nature Reviews Genetics (2024). 10.1038/s41576-023-00676-9

4 Murtazalieva, K., Mu, A., Petrovskaya, A. & Finn, R. D. The growing repertoire of phage anti-defence systems. Trends in Microbiology (2024). 10.1016/j.tim.2024.05.005

5 Amitsur, M., Levitz, R. & Kaufmann, G. Bacteriophage T4 anticodon nuclease, polynucleotide kinase and RNA ligase reprocess the host lysine tRNA. EMBO J 6, 2499–2503 (1987).

6 Loenen, W. A., Dryden, D. T., Raleigh, E. A., Wilson, G. G. & Murray, N. E. Highlights of the DNA cutters: a short history of the restriction enzymes. Nucleic Acids Res 42, 3–19 (2014). 10.1093/nar/gkt990

7 Koonin, E. V., Makarova, K. S. & Zhang, F. Diversity, classification and evolution of CRISPR-Cas systems. Curr Opin Microbiol 37, 67–78 (2017). 10.1016/j.mib.2017.05.008

8 Wein, T. & Sorek, R. Bacterial origins of human cell-autonomous innate immune mechanisms. Nature Reviews Immunology (2022). 10.1038/s41577-022-00705-4

9 Payne, L. J. et al. PADLOC: a web server for the identification of antiviral defence systems in microbial genomes. Nucleic Acids Research 50, W541–W550 (2022). 10.1093/nar/gkac400

10 Tesson, F. et al. Systematic and quantitative view of the antiviral arsenal of prokaryotes. Nature Communications 13, 2561 (2022). 10.1038/s41467-022-30269-9

11 DeWeirdt, P. C., Mahoney, E. M. & Laub, M. T. DefensePredictor: A Machine Learning Model to Discover Novel Prokaryotic Immune Systems. bioRxiv, 2025.2001.2008.631726 (2025). 10.1101/2025.01.08.631726

12 Mordret, E. et al. Protein and genomic language models chart a vast landscape of antiphage defenses. bioRxiv, 2025.2001.2008.631966 (2025). 10.1101/2025.01.08.631966

13 Uyttebroek, S. et al. Safety and efficacy of phage therapy in difficult-to-treat infections: a systematic review. The Lancet Infectious Diseases 22, e208–e220 (2022). 10.1016/S1473-3099(21)00612-5

14 Pirnay, J.-P. et al. Personalized bacteriophage therapy outcomes for 100 consecutive cases: a multicentre, multinational, retrospective observational study. Nature Microbiology 9, 1434–1453 (2024). 10.1038/s41564-024-01705-x

15 Brödel, A. K. et al. In situ targeted base editing of bacteria in the mouse gut. Nature 632, 877–884 (2024). 10.1038/s41586-024-07681-w

16 Forsberg, K. J. et al. Functional metagenomics-guided discovery of potent Cas9 inhibitors in the human microbiome. Elife 8 (2019). 10.7554/eLife.46540

17 Vassallo, C. N., Doering, C. R., Littlehale, M. L., Teodoro, G. I. C. & Laub, M. T. A functional selection reveals previously undetected anti-phage defence systems in the E. coli pangenome. Nature Microbiology 7, 1568–1579 (2022). 10.1038/s41564-022-01219-4

18 Hossain, A. A. et al. DNA glycosylases provide antiviral defence in prokaryotes. Nature (2024). 10.1038/s41586-024-07329-9

19 Forsberg, K. J. et al. Bacterial phylogeny structures soil resistomes across habitats. Nature 509, 612–616 (2014). 10.1038/nature13377

20 Clemente, J. C. et al. The microbiome of uncontacted Amerindians. Sci Adv 1 (2015). 10.1126/sciadv.1500183

21 Pehrsson, E. C. et al. Interconnected microbiomes and resistomes in low-income human habitats. Nature 533, 212–216 (2016). 10.1038/nature17672

22 Fierer, N. et al. Comparative metagenomic, phylogenetic and physiological analyses of soil microbial communities across nitrogen gradients. ISME J 6, 1007–1017 (2012). 10.1038/ismej.2011.159

23 Gao, L. et al. Diverse enzymatic activities mediate antiviral immunity in prokaryotes. Science 369, 1077–1084 (2020). 10.1126/science.aba0372

24 Koonin, E. V., Makarova, K. S., Wolf, Y. I. & Krupovic, M. Evolutionary entanglement of mobile genetic elements and host defence systems: guns for hire. Nat Rev Genet 21, 119–131 (2020). 10.1038/s41576-019-0172-9

25 Makarova, K. S., Wolf, Y. I., Snir, S. & Koonin, E. V. Defense islands in bacterial and archaeal genomes and prediction of novel defense systems. J Bacteriol 193, 6039–6056 (2011). 10.1128/JB.05535-11

26 Bateman, A. et al. The Pfam protein families database. Nucleic Acids Res 28, 263–266 (2000).

27 Haft, D. H. et al. TIGRFAMs: a protein family resource for the functional identification of proteins. Nucleic Acids Res 29, 41–43 (2001).

28 Payne, L. J., Hughes, T. C. D., Fineran, P. C. & Jackson, S. A. New antiviral defences are genetically embedded within prokaryotic immune systems. bioRxiv, 2024.2001.2029.577857 (2024). 10.1101/2024.01.29.577857

29 Lehman, I. R. & Pratt, E. A. On the Structure of the Glucosylated Hydroxymethylcytosine Nucleotides of Coliphages T2, T4, and T6. Journal of Biological Chemistry 235, 3254–3259 (1960). 10.1016/S0021-9258(20)81347-7

30 Wilson, G. G., Young, K. Y., Edlin, G. J. & Konigsberg, W. High-frequency generalised transduction by bacteriophage T4. Nature 280, 80–82 (1979). 10.1038/280080a0

31 Bryson, A. L. et al. Covalent Modification of Bacteriophage T4 DNA Inhibits CRISPR-Cas9. mBio 6, e00648 (2015). 10.1128/mBio.00648-15

32 Niu, Y., Suzuki, H., Hosford, C. J., Walz, T. & Chappie, J. S. Structural asymmetry governs the assembly and GTPase activity of McrBC restriction complexes. Nature Communications 11, 5907 (2020). 10.1038/s41467-020-19735-4

33 Swinton, D. et al. Purification and characterization of the unusual deoxynucleoside, alpha-N-(9-beta-D-2’-deoxyribofuranosylpurin-6-yl)glycinamide, specified by the phage Mu modification function. Proceedings of the National Academy of Sciences 80, 7400–7404 (1983). 10.1073/pnas.80.24.7400

34 Silva, R. M. B. et al. Modification of DNA by a viral enzyme and charged tRNA. bioRxiv, 2023.2003.2024.534169 (2023). 10.1101/2023.03.24.534169

35 Weigel, C. & Seitz, H. Bacteriophage replication modules. FEMS Microbiol Rev 30, 321–381 (2006). 10.1111/j.1574-6976.2006.00015.x

36 Maffei, E. et al. Systematic exploration of Escherichia coli phage–host interactions with the BASEL phage collection. PLOS Biology 19, e3001424 (2021). 10.1371/journal.pbio.3001424

37 Kemp, P., Garcia, L. R. & Molineux, I. J. Changes in bacteriophage T7 virion structure at the initiation of infection. Virology 340, 307–317 (2005). 10.1016/j.virol.2005.06.039

38 Hu, B., Margolin, W., Molineux, I. J. & Liu, J. The Bacteriophage T7 Virion Undergoes Extensive Structural Remodeling During Infection. Science 339, 576–579 (2013). 10.1126/science.1231887

39 Zheng, J. et al. Conformational changes in and translocation of small proteins: insights into the ejection mechanism of podophages. Journal of Virology 0, e01249–01224 (2024). 10.1128/jvi.01249-24

40 Vogt, J. & Schulz, G. E. The structure of the outer membrane protein OmpX from Escherichia coli reveals possible mechanisms of virulence. Structure 7, 1301–1309 (1999). 10.1016/S0969-2126(00)80063-5

41 Vica Pacheco, S., García González, O. & Paniagua Contreras, G. L. The lom gene of bacteriophage lambda is involved in Escherichia coli K12 adhesion to human buccal epithelial cells. FEMS Microbiol Lett 156, 129–132 (1997). 10.1111/j.1574-6968.1997.tb12717.x

42 Liu, X., Jiang, H., Gu, Z. & Roberts, J. W. High-resolution view of bacteriophage lambda gene expression by ribosome profiling. Proc Natl Acad Sci U S A 110, 11928–11933 (2013). 10.1073/pnas.1309739110

43 Dedrick, R. M. et al. Prophage-mediated defence against viral attack and viral counter-defence. Nat Microbiol 2, 16251 (2017). 10.1038/nmicrobiol.2016.251

44 Getz, L. J. & Maxwell, K. L. Diverse Antiphage Defenses Are Widespread Among Prophages and Mobile Genetic Elements. Annual Review of Virology 11, 343–362 (2024). 10.1146/annurev-virology-100422-125123

45 Wang, Y. et al. Genetic manipulation of Patescibacteria provides mechanistic insights into microbial dark matter and the epibiotic lifestyle. Cell 186, 4803-4817.e4813 (2023). 10.1016/j.cell.2023.08.017

46 Gios, E., Mosley, O. E., Takeuchi, N. & Handley, K. M. Genetic exchange shapes ultra-small Patescibacteria metabolic capacities in the terrestrial subsurface. bioRxiv, 2022.2010.2005.510940 (2022). 10.1101/2022.10.05.510940

47 Tian, R. et al. Small and mighty: adaptation of superphylum Patescibacteria to groundwater environment drives their genome simplicity. Microbiome 8, 51 (2020). 10.1186/s40168-020-00825-w

48 Bouderka, F. et al. Culture- and genome-based characterization of a tripartite interaction between patescibacterial epibionts, methylotrophic proteobacteria, and a jumbo phage in freshwater ecosystems. bioRxiv, 2024.2003.2008.584096 (2024). 10.1101/2024.03.08.584096

49 Burmeister, A. R. et al. Pleiotropy complicates a trade-off between phage resistance and antibiotic resistance. Proceedings of the National Academy of Sciences 117, 11207–11216 (2020). 10.1073/pnas.1919888117

50 Bailone, A. & Devoret, R. Isolation of ultravirulent mutants of phage λ. Virology 84, 547–550 (1978). 10.1016/0042-6822(78)90273-8

51 Abedon, S. T. Bacteriophage Adsorption: Likelihood of Virion Encounter with Bacteria and Other Factors Affecting Rates. Antibiotics 12 (2023).

52 Machnicka, M. A., Kaminska, K. H., Dunin-Horkawicz, S. & Bujnicki, J. M. Phylogenomics and sequence-structure-function relationships in the GmrSD family of Type IV restriction enzymes. BMC Bioinformatics 16, 336 (2015). 10.1186/s12859-015-0773-z

53 Van Valen, L. A New Evolutionary Law. Evol Theory (1973).

