## Supplementary Methods for "Metagenomic selections reveal diverse antiphage defenses in human and environmental microbiomes"

### Material and Methods

#### Metagenomic libraries

Metagenomic libraries, described previously, were generously shared by Gautam Dantas from Washington University in St. Louis. These libraries were constructed using oral swabs from Yanomani Amerindians[1], fecal samples from periurban residents of Lima, Peru[2] or soil samples collected from grasslands in the Cedar Creek Ecosystem Science Reserve (CC; Bethel, MN, USA)[3,4]. These libraries comprised 1.4-9.0 x $10^6$ unique clones with an average DNA insert size of 2 Kb (Table S1). All libraries were previously constructed in a pZE21 MCS1 plasmid backbone marked by kanamycin resistance ($Kan^R$) and were transformed into ultra-competent *E. coli* (MegaX DH10B $T1^R$ Electrocomp™ Cells; ThermoFisher cat. #C640003)[1-3]. The processing and storage of each metagenomic library was performed as previously described[5].

#### Metagenomic functional selection for phage-defense systems

Historical and recent work has established that intermediate phage concentrations in soft agar overlays can recover both direct phage defense mechanisms (which block infection without harming the infected bacterial cell) and abortive infection-based defenses (which trigger the death or dormancy of an infected cell to prevent phage replication and protect uninfected cells in the population)[6,7]. Thus, for each phage, we tested a series of phage concentrations to identify the minimal multiplicity of infection (MOI) at which an empty vector control was maximally eliminated. Due to different infectivity strengths among our panel of phages tested, different MOIs were used in selections with different phages (see Table S4 for details).

For the first iteration of the functional selection, vials of freezer stock from each library were thawed on ice and a volume of cells corresponding to approximately 10 times the number of unique clones in the library transferred to a microcentrifuge tube (Table S4). A similar number of empty vector control cells were used in parallel infections that contained the pZE21 cloning vector lacking a DNA insert. Cells were pelleted by centrifugation at 10,000 rpm for 1 minute to remove the supernatant and resuspended in a volume of LB broth (10 g/L casein peptone, 10 g/L NaCl, 5 g/L ultra-filtered yeast powder) with 50 ug/ml kanamycin (Kan) between ~10µl and 1ml. Resuspension volumes were chosen to achieve a 1:1 volumetric ratio with the amount of phages added to the solution, which were stored and diluted in SM buffer (100mM NaCl, 8mM $MgSO_4$, 50mM Tris-Cl, pH 7.5). Cells and phages were mixed and incubated at 37°C for 15 minutes to allow phage adhesion. After the incubation period, 3 ml of top agar (25 g/L of LB Broth + 5 g/L of Agar Granulated; *Fisher BioReagents* BP9724-500) was added to the bacteria phage mixture and poured onto approximately 15 mL of pre-hardened LB-Agar (10 g/L casein peptone, 10 g/L NaCl, 5 g/L ultra-filtered yeast powder, 12 g/L Agar) plates with 50 ug/ml Kan. Once the top agar had solidified, plates were incubated at 37°C overnight. The surviving colonies from this first round of selection were collected in 3 ml of LB-Broth by scraping them with a sterile L-shaped cell scraper (Fisher Scientific cat. #03-392-151) to gently remove them from the top agar (see Table S4 for surviving colony counts). One-third of the cells were used to make -80°C freezer stocks, and plasmids from the remaining 2 ml were purified using the ZymoPURE™ Plasmid Miniprep Kit (cat #D4210) using manufacturer recommendations. These plasmid preparations were used in subsequent selection experiments. Some selections with phage T4 were not collected by scraping and individual colonies were instead picked and processed as is described below.

For phages other than T4, a second iteration of selection was performed. In these instances, the isolated plasmids from colonies surviving the first round of phage infection were electroporated into ultra-competent *E. coli* MegaX. As a control, 100 ng of pZE21 vector expressing KanR gene was also transformed. All the electroporations were performed in a 0.1 cm gene pulser cuvette (Bio-Rad; cat.

#165-2089) using a Bio-Rad Gene Pulser Xcell with the following settings: 2.0 kV, 200 Ohms, 25 $\mu$F. Following electroporation, cells recovered for two hours at 37°C in Recovery Medium (Invitrogen™; cat. #C640003). A small volume of cells were used to determine the colony forming units (CFU) post-recovery. The remaining cells were inoculated into 25 ml LB broth with 50 ug/ml Kan and grown at 37°C until reaching an OD600 value of ~ 0.7, mixed with 70% LB-Glycerol, and flash-frozen. A small volume of frozen cells was thawed and used to determine a post-expansion, post-freeze titer for use in downstream experiments. Three selections using phage T7 repeatedly failed to transform well at this stage and were removed from further consideration. In all other cases, a second round of phage infection was performed exactly as occurred in iteration #1, with the following modifications. First, the number of phage particles in each $2^{nd}$-round selection was held at a near-constant level for a given phage (roughly 1-8x$10^8$ PFU/selection, depending on the phage). Second, each *E. coli* library was serially diluted 10-fold and each dilution plated with the same number phage particles (see Table S5 for details). Diluting libraries allowed us to identify plates with distinct phage-resistant colonies, as opposed to lawns of phage resistant cells (which often occurred, as some once-selected libraries were replete with phage resistance). A downside of using diluted libraries is a potential bottleneck in the data, which means that our reported phage resistance should be interpreted as a lower bound on the number of true resistance determinants in each library. This downside is outweighed by the high-confidence dataset that results from our approach. Using dilutions with distinct phage-resistant colonies increases the probability of recovering *bona-fide* resistance genotypes, as opposed to potential false positives that withstand infection due to their proximity to nearby clones with phage-eliminating properties.

### Sample preparation and sequencing

We sequenced plasmids from 60 of the 63 selections described above, omitting only the three T7 selections excluded previously. For eight selections with phage T4 (all except library O5), colonies were picked individually rather than collected into pools. From these selections, 66 colonies were grown overnight in LB-Kan, plasmids purified by ZymoPURE™ Miniprep (cat #D4210), and sequenced individually using Plasmidsaurus whole plasmid sequencing. These 66 plasmid preparations encoded 31 unique metagenomic DNA inserts listed in Table S6, 30 of which exhibited phage resistance (uniqueness is defined as sequences with less than 98% nucleotide identity over 98% the length of the shorter fragment). To aid with the subsequent analysis of pooled metagenomic plasmid preparations, we used a subset of these sequenced plasmids to generate a mock functional selection, comprised solely of known plasmids with confirmed phage resistance. This positive control plasmid mixture contained 13 plasmids encoding DNA inserts from 10 sequence groups. In seven instances, metagenomic plasmids with unrelated DNA inserts were added to the pool. In the three other cases, two plasmids with very similar insert sequences were pre-mixed and added to the pool together (nucleotide identity in each pair ranged from 93.5% to 100% over the length of the shorter sequence). Because we observed multiple instances of related but distinct insert sequences among our set of 66 picked colonies, these mixtures were intended to simulate the true diversity present in a mixed plasmid pool from one of our functional selections. Each of the 10 sequence groups was added to the final mixture at varying molar abundances, ranging from 3.0% to 17.1% of the total DNA in solution. As we outline later, this defined mixture helped us to confidently assign quality thresholds when assembling and curating data from pooled functional selections.

From the remaining 52 selections, colonies were collected and plasmids purified in aggregate via ZymoPURE miniprep, as described above. Pooled plasmid mixtures were sent to Plasmidsaurus for direct sequencing with Oxford Nanopore long-read technology (v14 chemistry, guppy base calling with super-accurate mode). Raw reads were filtered for quality and length using the program NanoFilt (v2.8.0) from the NanoComp package with parameters '-q 10 -l 500' (phred quality scores ≥10 and length ≥ 500 bp)[8]. Filtered reads were then used to assemble the metagenomic DNA inserts encoded by twice-selected libraries.

**Assembly, quality control, and annotation of metagenomic DNA inserts**

Filtered reads were converted to fasta format and bases originating from the pZE21 cloning vector were identified using the program cross_match (v1.090518) with the following parameters: -minmatch 12 -minscore 20 -screen[9]. Blocks of non-vector sequences longer than 50bp were retained and assembled into initial contigs using Flye (v2.9.2, parameters: --nano-hq, --meta, --scaffold), a long-read assembly algorithm designed for reads generated with Oxford Nanopore Technology[10]. To determine the cloned orientation of a DNA insert in pZE21 and to refine the contig ends near this vector junction, a custom script was used. This script considered reads with at least 25 perfectly matching bases to both the pZE21 cloning junction (its HincII site) and the end of an assembled contig. For each contig, the orientation of the cloned DNA insert was assigned by majority rule after counting reads that support either the forward or reverse orientation. These reads were then examined to assess whether any nucleotides were present between the vector junction and the end of each assembled contig, to potentially identify over-trimmed assemblies. In some cases, additional sequence was identified from these reads. These sequences were extracted for each contig, and aligned using MUSCLE (v3.8.31) with just one non-default parameter (-maxiters 2)[11]. From these alignments, consensus sequences were built for each new contig end with CIAlign (v1.1.0) and the parameter '--make_consensus'[12]. These consensus end sequences were then appended to their source contig in the appropriate orientation. Contigs with these appendages are indicated by a lower case 'a' at the end of each contig name. In some cases, coverage over these ends was examined by eye in the Geneious Prime sequence analysis suite (v2022.2.2) and manually trimmed based on dramatic drops in coverage (these contigs were renamed with a lower case 'm' suffix).

To determine the coverage of each assembled sequence, vector-cleaned reads were mapped to each contig using minimap2 (v2.24) with the parameters -ax map-ont[13]. For each contig, we normalized the 'mean depth of coverage' statistic by sample read number (normalized per 1,000 reads) and then log-transformed this normalized value (Table S6, Figure S8). From our mock functional selection (comprised of only previously validated phage defense plasmids), 13 of the 14 most abundant contigs mapped with high confidence to the known DNA sequence spiked into this control sample. The log-normalized coverage values for these 13 contigs ranged from 0.776 to 2.071 and correlated well with the molar equivalents added for each of the spiked-in sequence groups (Figure S8). The least abundant control plasmid was spiked into the defined mixture at 3.0% of total DNA and its corresponding contig had a coverage value of 0.776. Thus, we applied a minimum coverage filter of 0.775 to our full dataset, which we estimate corresponds to a DNA insert at roughly 3% relative abundance in its input plasmid pool. We also discarded contigs longer than 10 Kb and which had a raw 'mean depth of coverage' statistic less than 5. These filters helped to prioritize successful phage defense inserts and eliminate potential contaminating sequences, which were more common among long and low coverage contigs, based on our examination of the control plasmid mixture.

One high-coverage contig was assembled from our defined plasmid control which did not map to any input sequence (coverage value = 0.8). Thus, we manually examined all remaining contigs for obvious errors or sources of contamination. In aggregate, this check revealed 23 sequences from the *E. coli* MG1655 genome (present in all phage-resistant colonies and a common contaminant in molecular biology reagents), 11 sequences matching the human genome (which may have been introduced during sample processing), and 4 low-complexity sequences (potentially representing assembly artefacts)[14,15]. Blastn was used to identify contigs with more than 90% nucleotide identity to the *E. coli* MG1655 and human genomes (over 50% and 25% of contig length, respectively) and Dustmasker (v1.0.0) was used with default parameters to identify contigs for which >10% of their sequence was low-complexity DNA[16]. After removing these 38 contigs, 172 assembled contigs remained across the 52 selections subjected to pooled plasmid sequencing. As described above, individual colonies were also picked from eight T4 selections to yield 31 unique DNA sequences. These sequences were combined to yield a final dataset of 203 putative phage defense inserts from 60 independent phage selections (Table S6, Supplemental Dataset 1).

Defense inserts were annotated as follows. We first used the gene-finder MetaGeneMark to predict ORFs using default parameters[17]. To ascribe general functions to these ORFs, we used their amino acid sequences in a profile HMM search with HMMER3 against TIGRFAM and Pfam profile HMM databases[18-20]. The highest scoring profile was used to tentatively annotate each ORF. To facilitate visual inspection of these annotations, we erred on the side of promiscuously assigning general functions and so did not use an e-value cutoff for these HMMs. Thus, these generic annotations should be treated with some caution. In contrast, took a more conservative approach for annotating phage defense genes, yielding higher quality, but less encompassing, predictions. As before, we used HMMER3 to compare proteins sequences against HMM databases, this time scanning both DefenseFinder (v1.1) and Padloc2 (v2.0) phage defense profiles[21,22]. Only hits with an e-value better than $1e^{-5}$ were used to annotate ORFs, with the best-scoring hit being used if both databases yielded high quality matches.

### Comparisons to NCBI databases

Blastn was used to map defense inserts to putative defense islands in NCBI genomes. 203 defense inserts were compared against NCBI's nt database on October 24, 2024 and hits with over 70% nucleotide identity spanning at least 50% of the query sequence were considered. Of the initial 203 queries, 106 inserts had at least one blast hit that cleared these thresholds. NCBI sequences for up to 10 blast hits from each of these 106 queries were then downloaded, prioritizing those with the best e-value. The sequences +/- 10 Kb from the bounds of the blast hit were then extracted and annotated with DefenseFinder and Padloc2, as described above[21,22]. Of the 106 inserts subjected to this analysis, 99 mapped to a sequence with a nearby ORF that could be confidently assigned a phage defense function (Table S8).

Bacterial homologs of the seven ORFs depicted in Figure 5A were identified among NCBI proteins. Amino acid sequences from each predicted ORF were compared against NCBI's nr database using a blastp search performed on January 8$^{th}$, 2025. We defined homologs as hits with e-value better than $1e^{-3}$ and which had more than 35% amino acid identity over at least half of the query length. For each homolog, the NCBI protein ID was used to retrieve taxonomic information via NCBI's E-utilities interface. These taxonomic tables were then subjected to light manual curation to ensure consistent taxonomic nomenclature. For query each protein, homolog counts were cataloged by phylum and then normalized the number of proteins (in millions) for that phylum, as listed in the NCBI taxonomy database on January 10$^{th}$, 2025. Log-transformed values for these normalized counts are depicted in Figure 5C.

### Phylogenetic analyses

The phylogenetic origins for each of the 203 phage defense inserts were predicted by comparing their sequences to the GTDB database (v214.1) using the 'easy-taxonomy' workflow within the mmseqs2 software package (version: bb0a1b3569b9fe115f3bf63e5ba1da234748de23)[23,24]. AnnoTree v2.0 was used visualize these phyla on a bacterial genome tree, with each tip representing a bacterial class and identified phyla colored differently[25]. The tree was cropped at a defined radius. Previously published 16S rDNA sequencing experiments from our fecal, oral, and soil metagenomes were also analyzed to independently assess the bacterial community composition of these samples. For fecal and soil samples, operational taxonomic unit (OTU) tables were downloaded from previous publications[2,4]. For oral samples, OTU tables were not available and so the original 16S rDNA reads were downloaded and reanalyzed using the Qiime2 (release qiime2-2023.7) standard workflow (ENA accessions ERR687991, ERR687994, ERR687988 correspond to samples O3, O5, and O23, respectively)[1,26]. Briefly, reads were imported and quality-filtered using standard Qiime2 procedures (described in detail at docs.qiime2.org/2023.7/tutorials/overview). Reads were then denoised using the deblur method with a trim length of 115. To assign OTU taxonomy, reads were compared against the SILVA 138 full 16S pre-trained classification dataset released with Qiime2 and available at the website: docs.qiime2.org/2023.7/data-resources[27-29]. The detailed command is as follows: qiime feature-classifier classify-sklearn --i-classifier silva-138-99-nb-classifier.qza --i-reads rep-seqs.qza --o-classification taxonomy.qza. Barplots were then made using the ‘qiime taxa barplot’ command and visualized online using Qiime2 View (view.qiime2.org). The corresponding OTU table was then downloaded at the ‘class’ resolution and analyzed alongside fecal and soil OTU tables.

To enable comparisons against phage defense contigs, 16S OTU tables were first converted from NCBI taxonomic labels to a consistent GTDB-based taxonomic labeling scheme. This was important to ensure consistent labels for datasets generated independently and analyzed at different times. To achieve this relabeling, OTU tables for fecal, oral, and soil biomes were extracted at the resolution of family, class, and class, respectively. Each label (e.g. class Bacteroidia) was then used in a manual search against the GTDB online database (release 08-RS214) with the option 'GTDB species representatives only' flagged. The GTDB phylum that was most consistently associated with each NCBI class or family was then associated with that query. A support score was calculated for each taxonomic association by dividing the number of GTDB accessions in the search results with the assigned phylum by the total number of accessions returned. Of the 85 associations made across all datasets, 84 had a support score of 0.9 or higher. With consistent GTDB assignments in hand for all samples, comparisons between 16S and phage defense datasets were performed at the resolution of phylum. This high-level classification maximized the sample size under consideration (detailed taxonomic predictions are challenging for short metagenomic DNA sequences) and made our conclusions more robust to differences associated with 16S datasets generated at different times by different authors. For each of the nine metagenomes analyzed for phage defense and 16S composition, a phylum’s relative abundance in each dataset was tabulated and compared via linear regression in R (Figure S2).

Homologs of OmpA, OmpX, $OMP_{Tell\text{-}1}$, and $OMP_{Tell\text{-}2}$ were retrieved from NCBI via BlastP on September 12, 2024. They were aligned via muscle and a maximum likelihood phylogeny generated using PhyML with 100 bootstraps. Structural predictions were generated using Alphafold3 via the website alphafoldserver.com and signal peptides predicted using the SignalP 6.0 webserver[30,31].

### Cloning

To amplify metagenomic inserts from mixed plasmid pools following phage selection, we designed primers with homology to both the target insert and the pZE21 vector to facilitate Gibson Cloning (see Table S10 for primers and PCR conditions). Common inverse PCR primers were used to linearize pZE21. To generate point mutations in a contig or cloned open reading frame, primers with the designed variant were used with constant primers on the pZE21 backbone to generate two fragments with homologous ends to facilitate Gibson Cloning (see Table S10 for primer sequences, and detailed PCR parameters). Generically, all PCRs were performed with the Q5 DNA polymerase (High-Fidelity 2x Master Mix, NEB cat. #M0492S) for a total of 35 cycles according to manufacturer's recommendations. After PCR, amplicons were gel purified using the Zymoclean™ Gel DNA Recovery Kit (ZYMO, cat. #D4007) per suggested protocols and 1ul of each amplicon for a given cloning reaction was mixed with 2ul of NEB HiFi Assembly Master Mix (NEB, cat. #M5520AA) to give a 4ul total reaction that was incubated at 50ºC for 60 minutes. Subsequently, 1ul was used to transform 25ul of chemically competent *E. coli* MegaX cells and selected on kanamycin to isolate transformants. Individual colonies were picked, grown overnight shaking at 37ºC, plasmids purified, and sequence verified by whole plasmid sequencing (Plasmidsaurus) before downstream experimentation. In some cases, sequences were synthesized by GenScript and cloned into pZE21_tetR, swapped directly for the GFP gene in this construct (Table S11).

To facilitate protein purification, the coding sequences for MdgA and PD-T4-$3_{Capno}$ were cloned into the ppSumo vector which contains an N-terminal 6xHis-Sumo-tag. Amino acid mutations were introduced using the Agilent QuickChange site-directed mutagenesis kit. Sequences were confirmed by nanopore or Sanger sequencing.

**Plaque assays to measure phage defense**

To test phage defense, overnight cultures expressing an empty vector, a GFP control, or a putative defense were diluted 1:50 in LB-Broth+Kan50 supplemented with 1.5 mM MgCl2 and 1.5 mM CaCl2. These cultures were grown at 37°C until they reached an optical density (OD) range of 0.4-0.8. Subsequently, 100 µL of the bacterial outgrowth was mixed with 3 mL of 0.5% soft agar (6.25 g Fisher LB-Broth Miller Granulated, 1.25 g Fisher Agar Granulated, and 250 mL Millipore water) and poured onto an LB-agar plate containing 50 µg/mL kanamycin. A tenfold serial dilution of phage was prepared in SM buffer and, 2.5 µL of each dilution was spotted on the soft agar media. The plates were then incubated overnight at 37°C. To quantify phage protection, the efficiency of plaquing (EOP) was calculated relative to the GFP or empty vector control.

This assay was used to validate defense inserts, identify ORFs necessary for defense from these inserts, and test for ORF sufficiency. To test whether functional selections yielded *bona fide* phage defense, a set of 73 inserts were chosen for individual validation experiments. Inserts were isolated from single colonies following selection or by PCR amplifying them from post-selection plasmid mixtures, followed by cloning the insert amplicons into the pZE21-MCS1 plasmid. To identify the ORF required for phage protection, each intact ORF in a contig was individually mutated by deleting the start codon, using the cloning methods previously described. To test if an ORF or system was sufficient for phage defense, it was either cloned or synthesized into pZE21_TetR_GFP, swapped directly for the GFP gene in this vector. Plasmids, cloned sequences, and induction conditions are enumerated in table Sj. For consistency, all ORFs in Figure 5B were tested with inducer, though MdgA and MdgB were otherwise interrogated in non-inducing conditions given their strong defense phenotype, per table Sj. To induce expression, 100ng/mL doxycycline was used.

**Protein purification**

6xHis-Sumo-MdgA & 6xHis-Sumo-PD-T4-$3_{Capno}$ WT plasmids were transformed into *E. coli* Rosetta (DE3) competent cells for protein expression. Cells were inoculated into 5 mL LB starter cultures with the addition of kanamycin (50 µg/mL) and chloramphenicol (34 ug/mL) and grown overnight at 37 °C. Starter cultures were transferred to 1 L of LB with kanamycin (50 µg/mL) and chloramphenicol (34 ug/mL) and grown at 37 °C. Once the cells reached an OD600 of 0.5 to 0.8, they were induced with IPTG (0.4 mM) and grown overnight at 16 °C. The cells were then centrifuged at 4,000 x g for 15 min at 4 °C and resuspended in lysis buffer (300 mM NaCl, 50 mM Tris-HCl pH 8, 14.2 mM BME, and 1 mM PMSF). The cells were then lysed by sonication and cleared by centrifugation at 35,000 x g for 30 min at 4 °C. The cleared supernatants were incubated with Ni-NTA resin for 1 hr at 4 °C. Following incubation, the Ni-NTA resin was loaded onto a gravity-flow column, washed with 80 mL of wash buffer (50 mM Tris-HCl pH 8, 300 mM NaCl, 25 mM imidazole, and 5 mM BME), and eluted in elution buffer (50 mM Tris-HCl pH 8, 300 mM NaCl, 300 mM imidazole, and 5 mM BME) at 4 °C. The proteins were then incubated at °4 C overnight with Ulp1 protease to remove the 6xHis-Sumo-tag. The proteins were further purified by size-exclusion chromatography using a HiLoad 16/600 Superdex 200 pg column in gel filtration buffer (50 mM Tris-HCl pH 8, 300 mM NaCl, and 1 mM DTT). Peak fractions were pooled and concentrated using Amicon Ultra centrifugal filters. Purified proteins were aliquoted, flash frozen in liquid nitrogen, and stored at -80 °C.

Mutant proteins were purified as described above except that the temperature was lowered to 18 °C after induction, and a Superdex 200 Increase 10/300 GL column was used for the size-exclusion chromatography.

**Nuclease activity assays**

Phage genomic DNA was extracted from 10 to 15ml of high titer lysate using the Monarch HMW DNA Extraction Kit for Tissue (NEB #T3060) per manufacturer's recommendations for phage gDNA extraction except that proteinase K treatment was not performed. Genomic DNA and protein samples were quantified using a NanoDrop One spectrophotometer. For the reactions, 50 ng of DNA and the specified concentration of purified protein were incubated at 37 °C for 10 min in nuclease buffer (50 mM Tris-HCl pH 8, 1 mM $MgCl_2$, 1 mM $MnCl_2$) in a total volume of 40 μL. The reactions were stopped by the addition of 1 μL of 0.5 M EDTA pH 8 to each reaction. Reaction products were resolved by performing gel electrophoresis using 0.8% agarose gels containing ethidium bromide. The reaction products were then visualized using a ChemiDoc Imaging System.

**Isolation and sequencing of phage escaper mutants**

To isolate phage escaper mutants, we performed a solid-phage infection using the defense insert encoding $OMP_{Acin-4}$ (T7-4). We picked escaper phage plaques with a pipette tip into 50ul of SM buffer and carried out a tenfold serial dilution of these picked plaques on *E. coli* MegaX with an empty pZE21-MCS1 vector. We then again isolated three individual plaques in 50ul of SM buffer and added this volume to 100ul of mid-log T7-4 cells (OD 0.5-0.8) in soft agar overlays. We purified the resultant confluent lawn of plaques by resuspending it in 3ml of SM buffer and filtering through a 0.22 μM filter. Since a larger volume of phage stock was necessary for phage genomic DNA (gDNA) extraction, we conducted a second propagation in 50ml liquid media, again using the same background strain, as previously described[32]. As before, we used T7-4 encoding *E. coli* cells. The extraction of phage gDNA was carried out using the Monarch HMW DNA Extraction Kit for Tissue (NEB #T3060), and the extracted gDNA was sent for whole Illumina bacterial genome sequencing at SeqCenter with variants called by mapping against the wild-type T7 genome (NC_001604.1). Only one of three phage propagations yielded sufficient genomic DNA and read quality for high-confidence mapping.

**T7 phage absorption assay**

A T7 phage absorption assay was performed in a BW25113 background to take advantage of the *E. coli* knockout Keio collection[33]. As negative controls, we used BW25113 expressing pZE21-MCS1 and pZE21_TetR_GFP and used the Keio △WaaC mutant as a positive control. An overnight culture for each OMP and control strain was grown in 4mL LB-Kan50 supplemented with 5μL 1M $MgCl_2$, and 5μL 1M $CaCl_2$. The following day, 120uL of overnight culture was added to 6mL of LB-Kan50 supplemented with 7.5μL 1M $MgCl_2$, 7.5μL 1M $CaCl_2$ and 100 ng/ml doxycycline (to induce GFP or OMP expression, as indicated). The outgrowth was incubated at 37C while shaking at 200rpm until reaching an OD600 value between 0.4 and 0.7. Cells were then transferred to a 15mL conical tube and centrifugated at 4121rpm for 10 minutes. The supernatant was decanted, and cell pellet was resuspended in 1mL of LB-Broth. The resuspended cells were split into two microcentrifuge tubes (500 μL each) and T7 phages added to achieve an approximate MOI of 1 (two dilutions were used to ensure a close approximation of the desired MOI). MOIs were approximated using previously determined standard curves relating OD600 to CFU/ml values and estimates of phage T7 titers. Microcentrifuge tubes were then vortexed for 15 seconds and the phage-cell mixtures incubated using an Eppendorf Thermomixer at 37C, 300RPM for 5 minutes. Following incubation, the samples were centrifuged for 10 min at 4000 RCF. The phage supernatant was then carefully collected without disturbing the bacteria pellet in a sterile microcentrifuge tube. The T7 titers from these supernatants were determined by performing a standard plaque assay in solid media using BW25113/pZE21-MCS1 as cell background.

Phage titers required estimation because adsorption was optimal with very fresh phage preparations (less than three hours old). This did not leave time for *a priori* phage titering. Contemporaneous with adsorption, cultures and phage preparations were titered to empirically calculate actual MOIs the next day. Only experiments with empirical MOIs very close to one were considered (n=3 biological replicates). Free phages from supernatants are depicted as a proportion of input phages in Figure 4D, based on these input titer values.

**Liquid phage infections**

Overnight cultures were grown to mid-log in 4mL LB-Kan50 supplemented with 5µL 1M $MgCl_2$, and 5µL 1M $CaCl_2$ and then normalized to an $OD_{600}$ value of ~0.25 using this media. Where indicated in table Sj, doxycycline was added at 100ng/ml to this media to induce expression. For each diluted culture, 100µL of culture was added to a flat-bottom 96-well plate (Costar #3596). Phage dilutions were added to each well at an MOI of $10^{-2}$. SM buffer was added as negative control. Plates were incubated at 37C with orbital shaking 800rpm in a Bio Tek LogPhage600 plate reader. $OD_{600}$ was recorded every 10 minutes. Three technical replicates were conducted for each strain. When assaying for abortive infection phenotypes, it is important to quantify phage titers in experimental and control strains after OD values crash. To achieve this, we performed parallel infections from the same cultures, run simultaneously in a different 96-well plate. After observing a bacterial crash (~4 hours post-infection), 80µL from each well was collected from one plate in a microtube. Then, centrifugation at 16,000 rpm for 5 minutes was performed to precipitate the sample. Supernatant containing phages were collected and plaque assay performed to measure phage titers. The other plate was not touched such that a unperturbed growth curve could be observed (withdrawing 80µL would otherwise dramatically impact $OD_{600}$ readings).

References

1 Clemente, J. C. *et al.* The microbiome of uncontacted Amerindians. *Sci Adv* **1** (2015). https://doi.org:10.1126/sciadv.1500183
2 Pehrsson, E. C. *et al.* Interconnected microbiomes and resistomes in low-income human habitats. *Nature* **533**, 212-216 (2016). https://doi.org:10.1038/nature17672
3 Fierer, N. *et al.* Comparative metagenomic, phylogenetic and physiological analyses of soil microbial communities across nitrogen gradients. *ISME J* **6**, 1007-1017 (2012). https://doi.org:10.1038/ismej.2011.159
4 Forsberg, K. J. *et al.* Bacterial phylogeny structures soil resistomes across habitats. *Nature* **509**, 612-616 (2014). https://doi.org:10.1038/nature13377
5 Forsberg, K. J. *et al.* Functional metagenomics-guided discovery of potent Cas9 inhibitors in the human microbiome. *Elife* **8** (2019). https://doi.org:10.7554/eLife.46540
6 Takahashi, H., Coppo, A., Manzi, A., Martire, G. & Pulitzer, J. F. Design of a system of conditional lethal mutations (tab/k/com) affecting protein-protein interactions in bacteriophage T4-infected Escherichia coli. *Journal of Molecular Biology* **96**, 563-578 (1975). https://doi.org:https://doi.org/10.1016/0022-2836(75)90139-4
7 Vassallo, C. N., Doering, C. R., Littlehale, M. L., Teodoro, G. I. C. & Laub, M. T. A functional selection reveals previously undetected anti-phage defence systems in the E. coli pangenome. *Nature Microbiology* **7**, 1568-1579 (2022). https://doi.org:10.1038/s41564-022-01219-4
8 De Coster, W., D'Hert, S., Schultz, D. T., Cruts, M. & Van Broeckhoven, C. NanoPack: visualizing and processing long-read sequencing data. *Bioinformatics* **34**, 2666-2669 (2018). https://doi.org:10.1093/bioinformatics/bty149
9 Ewing, B., Hillier, L., Wendl, M. C. & Green, P. Base-calling of automated sequencer traces using phred. I. Accuracy assessment. *Genome Res* **8**, 175-185 (1998). https://doi.org:10.1101/gr.8.3.175
10 Kolmogorov, M., Yuan, J., Lin, Y. & Pevzner, P. A. Assembly of long, error-prone reads using repeat graphs. *Nature Biotechnology* **37**, 540-546 (2019). https://doi.org:10.1038/s41587-019-0072-8
11 Edgar, R. C. MUSCLE: multiple sequence alignment with high accuracy and high throughput. *Nucleic acids research* **32**, 1792-1797 (2004). https://doi.org:10.1093/nar/gkh340
12 Tumescheit, C., Firth, A. E. & Brown, K. CIAlign: A highly customisable command line tool to clean, interpret and visualise multiple sequence alignments. *PeerJ* **10**, e12983 (2022). https://doi.org:10.7717/peerj.12983
13 Li, H. New strategies to improve minimap2 alignment accuracy. *Bioinformatics* **37**, 4572-4574 (2021). https://doi.org:10.1093/bioinformatics/btab705
14 Salter, S. J. *et al.* Reagent and laboratory contamination can critically impact sequence-based microbiome analyses. *BMC Biology* **12**, 87 (2014). https://doi.org:10.1186/s12915-014-0087-z
15 Breitwieser, F. P., Pertea, M., Zimin, A. V. & Salzberg, S. L. Human contamination in bacterial genomes has created thousands of spurious proteins. *Genome Res* **29**, 954-960 (2019). https://doi.org:10.1101/gr.245373.118
16 Camacho, C. *et al.* BLAST+: architecture and applications. *BMC Bioinformatics* **10**, 421 (2009). https://doi.org:10.1186/1471-2105-10-421
17 Zhu, W., Lomsadze, A. & Borodovsky, M. Ab initio gene identification in metagenomic sequences. *Nucleic Acids Res* **38**, e132 (2010). https://doi.org:10.1093/nar/gkq275
18 Bateman, A. *et al.* The Pfam protein families database. *Nucleic Acids Res* **28**, 263-266 (2000).
19 Haft, D. H. *et al.* TIGRFAMs: a protein family resource for the functional identification of proteins. *Nucleic Acids Res* **29**, 41-43 (2001).
20 Finn, R. D., Clements, J. & Eddy, S. R. HMMER web server: interactive sequence similarity searching. *Nucleic Acids Res* **39**, W29-37 (2011). https://doi.org:10.1093/nar/gkr367
21 Tesson, F. *et al.* Systematic and quantitative view of the antiviral arsenal of prokaryotes. *Nature Communications* **13**, 2561 (2022). https://doi.org:10.1038/s41467-022-30269-9

22 Payne, L. J. *et al.* PADLOC: a web server for the identification of antiviral defence systems in microbial genomes. *Nucleic Acids Research* **50**, W541-W550 (2022). https://doi.org:10.1093/nar/gkac400
23 Steinegger, M. & Söding, J. MMseqs2 enables sensitive protein sequence searching for the analysis of massive data sets. *Nature Biotechnology* **35**, 1026-1028 (2017). https://doi.org:10.1038/nbt.3988
24 Parks, D. H. *et al.* A complete domain-to-species taxonomy for Bacteria and Archaea. *Nat Biotechnol* (2020). https://doi.org:10.1038/s41587-020-0501-8
25 Mendler, K. *et al.* AnnoTree: visualization and exploration of a functionally annotated microbial tree of life. *Nucleic Acids Research* **47**, 4442-4448 (2019). https://doi.org:10.1093/nar/gkz246
26 Bolyen, E. *et al.* Reproducible, interactive, scalable and extensible microbiome data science using QIIME 2. *Nature Biotechnology* **37**, 852-857 (2019). https://doi.org:10.1038/s41587-019-0209-9
27 Quast, C. *et al.* The SILVA ribosomal RNA gene database project: improved data processing and web-based tools. *Nucleic Acids Research* **41**, D590-D596 (2013). https://doi.org:10.1093/nar/gks1219
28 Robeson, M. S., II *et al.* RESCRIPt: Reproducible sequence taxonomy reference database management. *PLOS Computational Biology* **17**, e1009581 (2021). https://doi.org:10.1371/journal.pcbi.1009581
29 Bokulich, N. A. *et al.* Optimizing taxonomic classification of marker-gene amplicon sequences with QIIME 2's q2-feature-classifier plugin. *Microbiome* **6**, 90 (2018). https://doi.org:10.1186/s40168-018-0470-z
30 Teufel, F. *et al.* SignalP 6.0 predicts all five types of signal peptides using protein language models. *Nature Biotechnology* **40**, 1023-1025 (2022). https://doi.org:10.1038/s41587-021-01156-3
31 Abramson, J. *et al.* Accurate structure prediction of biomolecular interactions with AlphaFold 3. *Nature* **630**, 493-500 (2024). https://doi.org:10.1038/s41586-024-07487-w
32 Bonilla, N. *et al.* Phage on tap–a quick and efficient protocol for the preparation of bacteriophage laboratory stocks. *PeerJ* **4**, e2261 (2016). https://doi.org:10.7717/peerj.2261
33 Baba, T. *et al.* Construction of <i>Escherichia coli</i> K‐12 in‐frame, single‐gene knockout mutants: the Keio collection. *Molecular Systems Biology* **2**, 2006.0008 (2006). https://doi.org:https://doi.org/10.1038/msb4100050
