## Supplementary material for "Metagenomic selections reveal diverse antiphage defenses in human and environmental microbiomes": Figures S1 - S8

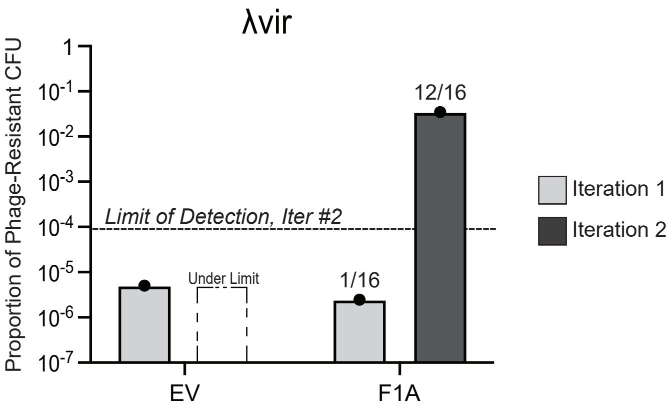


**Figure S1. A second round of phage selection enriches for plasmid-encoded defenses.** Proportion of phage resistance colonies for a control empty vector sample and the F1A library were subjected to two iterative λvir infections. Dashed lines represent the limit of detection for the second iteration, based on the number of cells plated. Fractions above the F1A bars indicate the proportion of isolated plasmids from each iteration that conferred λvir resistance over the number tested.

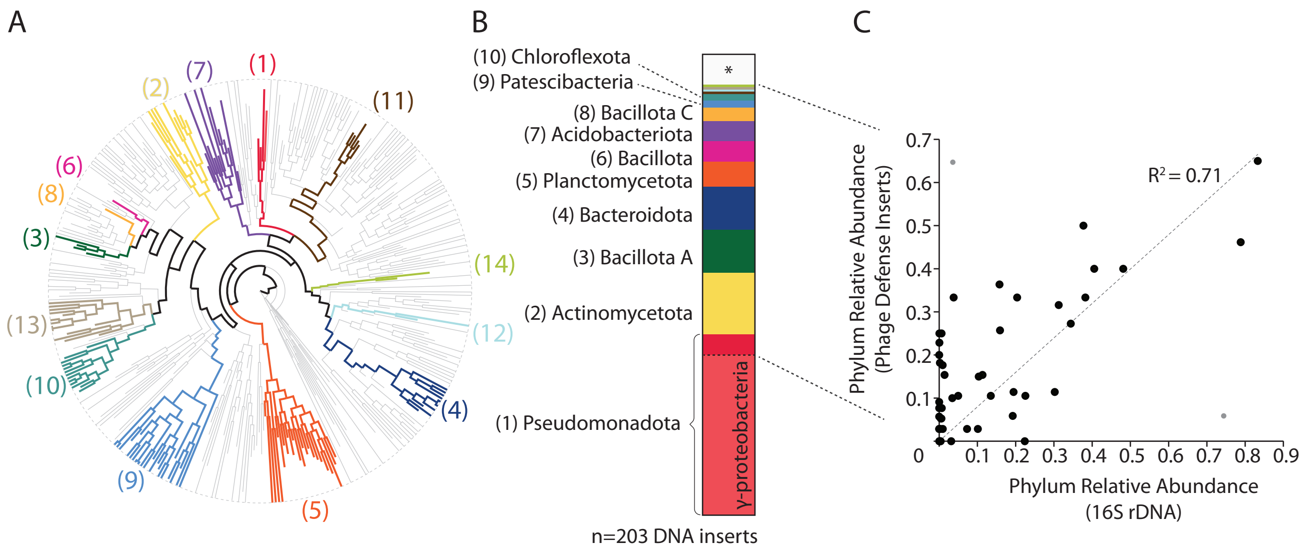


**Figure S2. A phylum's abundance in the phage defense dataset correlates with its abundance as measured by sequencing 16S rDNA amplicons.** (A, B) Reprinted from Figures 1F and 1G to give context for panel (C). (C) Regression analysis of a phylum's abundance in our phage defense dataset compared to its abundance as measured by 16S rDNA amplicon sequencing in its original fecal, oral, or soil metagenome. The γ-proteobacteria were omitted from both datasets before regression. Regression was performed with and without two outliers, depicted as gray points (Rosner's Extreme Studentized Deviate test, $p<0.005$). The outlier-excluded analysis is shown. If outliers are included, the correlation remains significant ($R^2=0.53$, $p=3e^{-9}$).

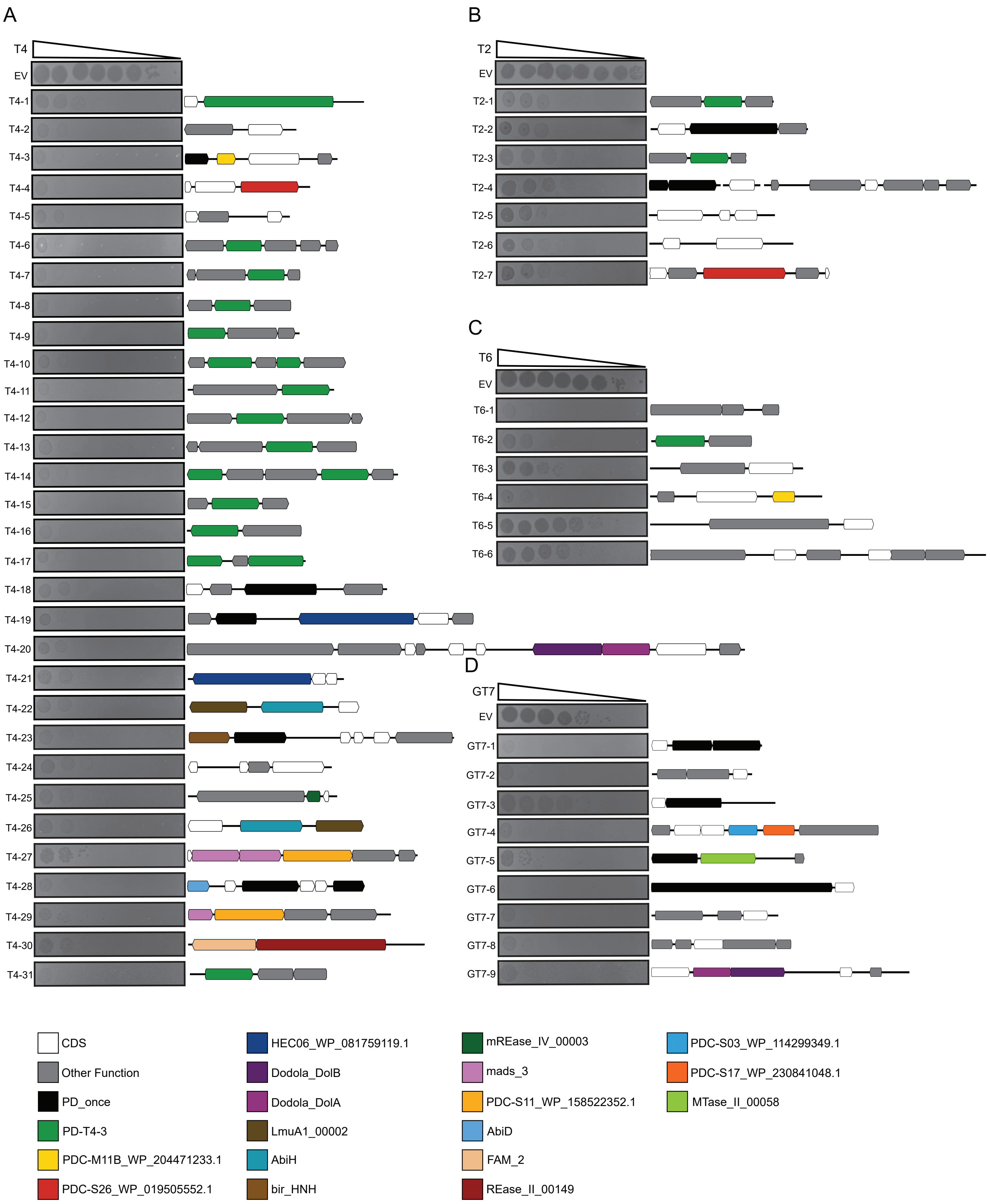


**Figure S3. Representative plaque assays for 64 validated DNA.**

(Figure continued on next page)

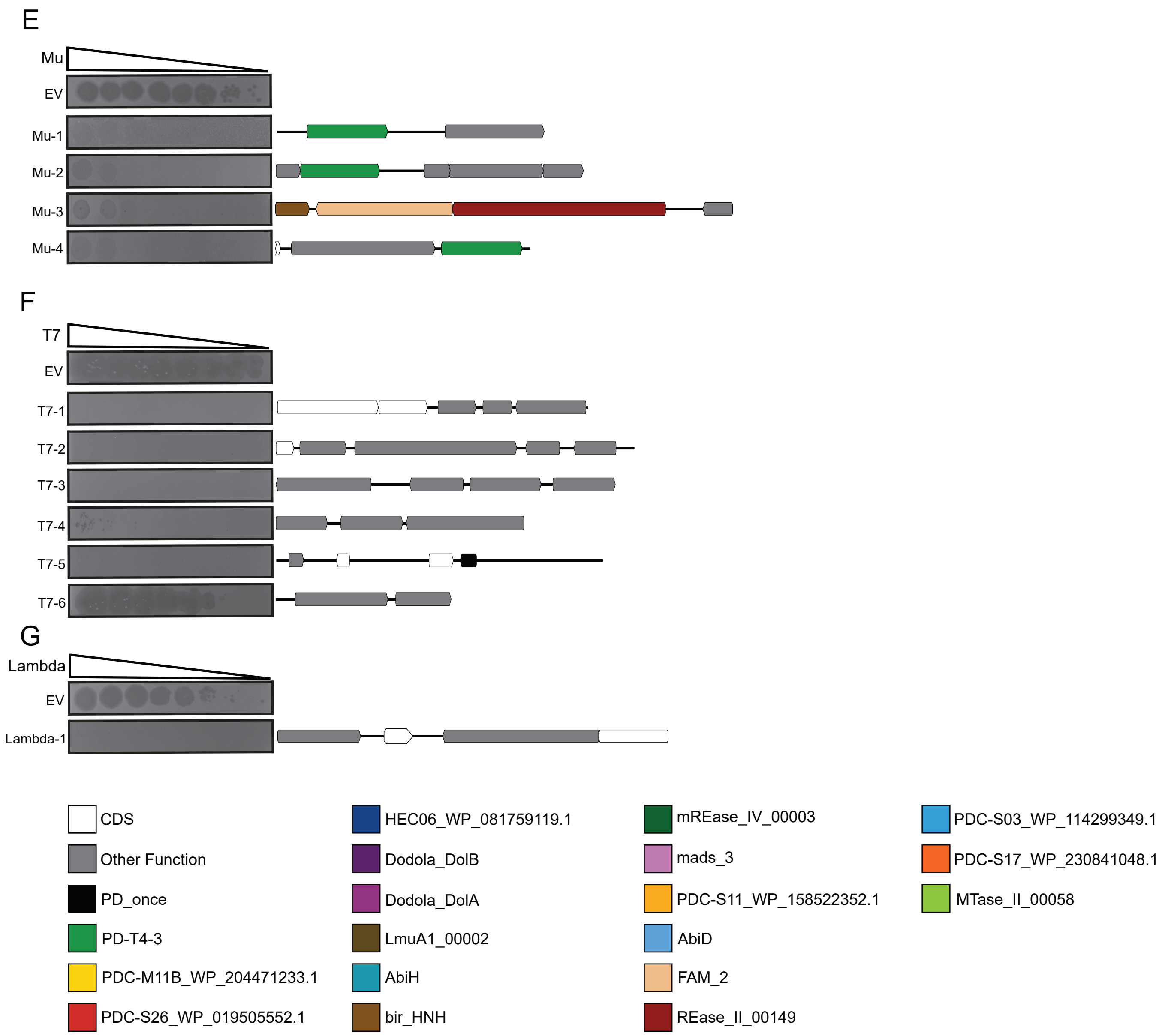


**Figure S3. Representative plaque assays for 64 validated DNA.** (A) Thirty-one validated DNA inserts from T4 selections. (B) Seven validated DNA inserts from T2 selections. (C) Six validated DNA inserts from T6 selections. (D) Nine validated DNA inserts from T4-GT7 selections. (E) Four validated DNA inserts from Mu selections. (F) Six validated DNA inserts from T7 selections. (G) One validated DNA insert from λvir selections. ORFs colored white represent genes of unknown function. ORFs colored grey indicate predicted functions without known links to phage defense. ORFs in various colors indicate matches to a Padloc2 or DefenseFinder domain profile with an e-value better than $1e^{-5}$. Colors match the legend below plaque images. The figure spans two pages.

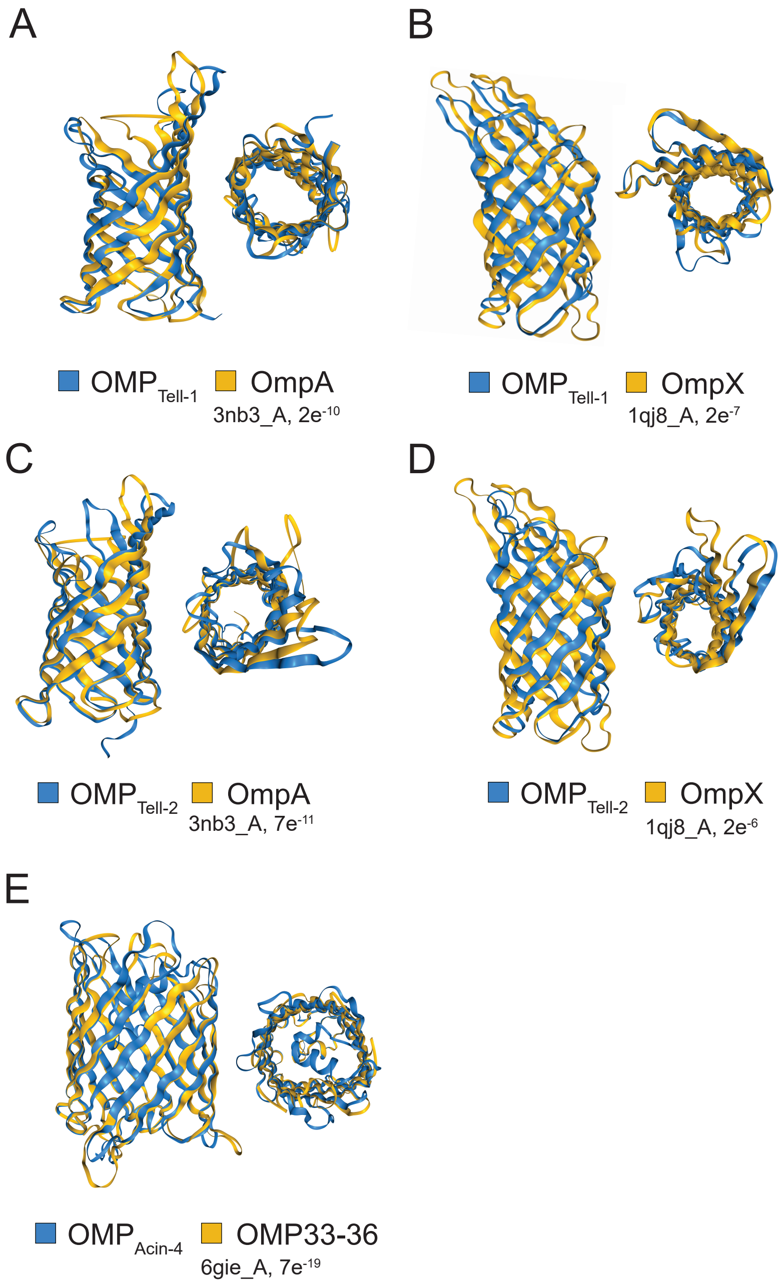


**Figure S4. Structural predictions by AlphaFold3 for $OMP_{Tell\text{-}1}$, $OMP_{Tell\text{-}2}$, and $OMP_{Acin\text{-}4}$ aligned with their respective structural homologs.** (A-B) $OMP_{Tell\text{-}1}$ predicted structure superimposed with (A) OMPA 3nb3_A and (B) OMPX 1qj8_A. (C-D) $OMP_{Tell\text{-}2}$ predicted structure superimposed with (C) OMPA 3nb3_A and (D) OMPX 1qj8_A. (E) $OMP_{Acin\text{-}4}$ predicted structure superimposed with OMP33-36 6gie_A. E-values are those reported by Foldseek to indicate the significance of a structural match. Images omit signal peptide sequences.

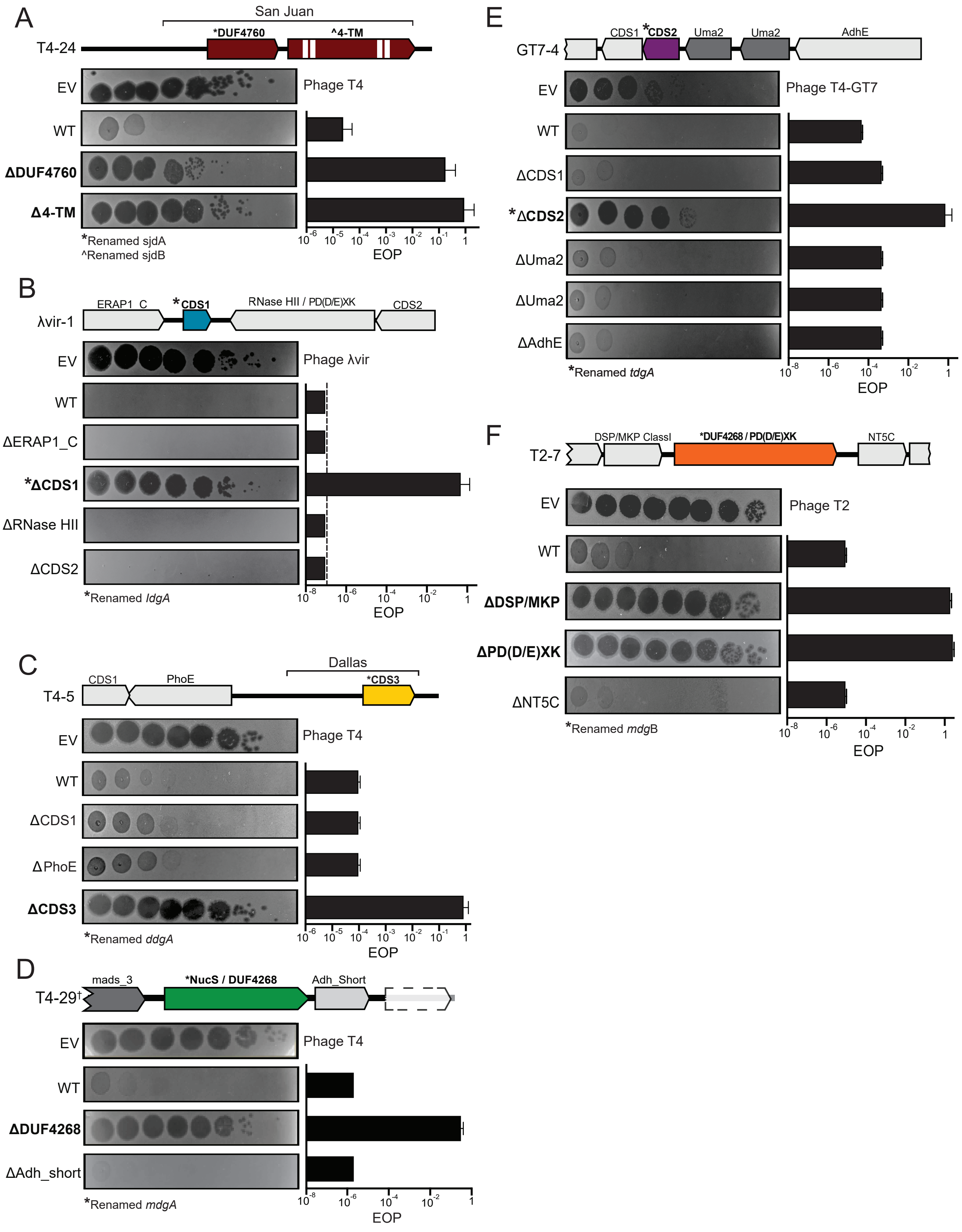


**Figure S5: Identifying gene(s) necessary for phage defense in unknown systems.**
Caption continues, next page.

**Figure S5: Identifying gene(s) necessary for phage defense in unknown systems.** Mutagenesis of seven defense inserts without known defense systems reveals gene(s) necessary for phage defense. To perturb each full-length ORF on each contig, the middle nucleotide of its predicted start codon was deleted. Empty vector and wild-type DNA inserts were included as controls. EOPs were calculated for each genotype relative to the empty vector control. Genes are colored if they are both necessary and sufficient for phage defense, either in isolation or as part of a defense system defined in brackets. Dark grey ORFs indicate defense-associate domains, per Figure S4. Other ORFs are colored in light grey. (A) A two-gene defense system is necessary and sufficient for defense against phage T4. A gene with a DUF4760 domain and another with four transmembrane domains (depicted in white) were renamed *sjdA* and *sjdB*, respectively. (B) A coding sequence (CDS) of unknown function is necessary and sufficient for defense against $\lambda_{vir}$, re-named *ldgA*. (C) A single-gene defense system is necessary and sufficient for defense against phage T4 (genetic sufficiency defined by the bracket). A CDS of unknown function was renamed *ddgA*. (D) A single ORF with NucS and DUF4268 domains is necessary and sufficient for defense against phage T4 and was renamed *mdgA*. The dagger symbol (†) indicates that a truncated variant of insert T4-29 was used in these experiments. Preliminary work indicated that this insert, generated accidentally during cloning, was sufficient for phage defense and lacked the ORF from T4-29 depicted in dashed lines. (E) A single CDS of unknown function is necessary and sufficient for defense against phage T4-GT7, renamed *tdgA*. (F) A single ORF with DUF4268 and PD(D/E)XK domains is necessary and sufficient for defense against phage T4, renamed *mdgB*. Another gene on the insert T2-7 also appears necessary for phage defense but was not sufficient for defense. Given that *mdgB* is sufficient to provide defense (Figure 5), we speculate that polar effects from deleting the stop codon in its upstream gene can result in the loss of phage defense in the T2-7 insert. All depicted experiments were performed in biological triplicate with representative plaque assays shown. Error bars depict standard deviation.

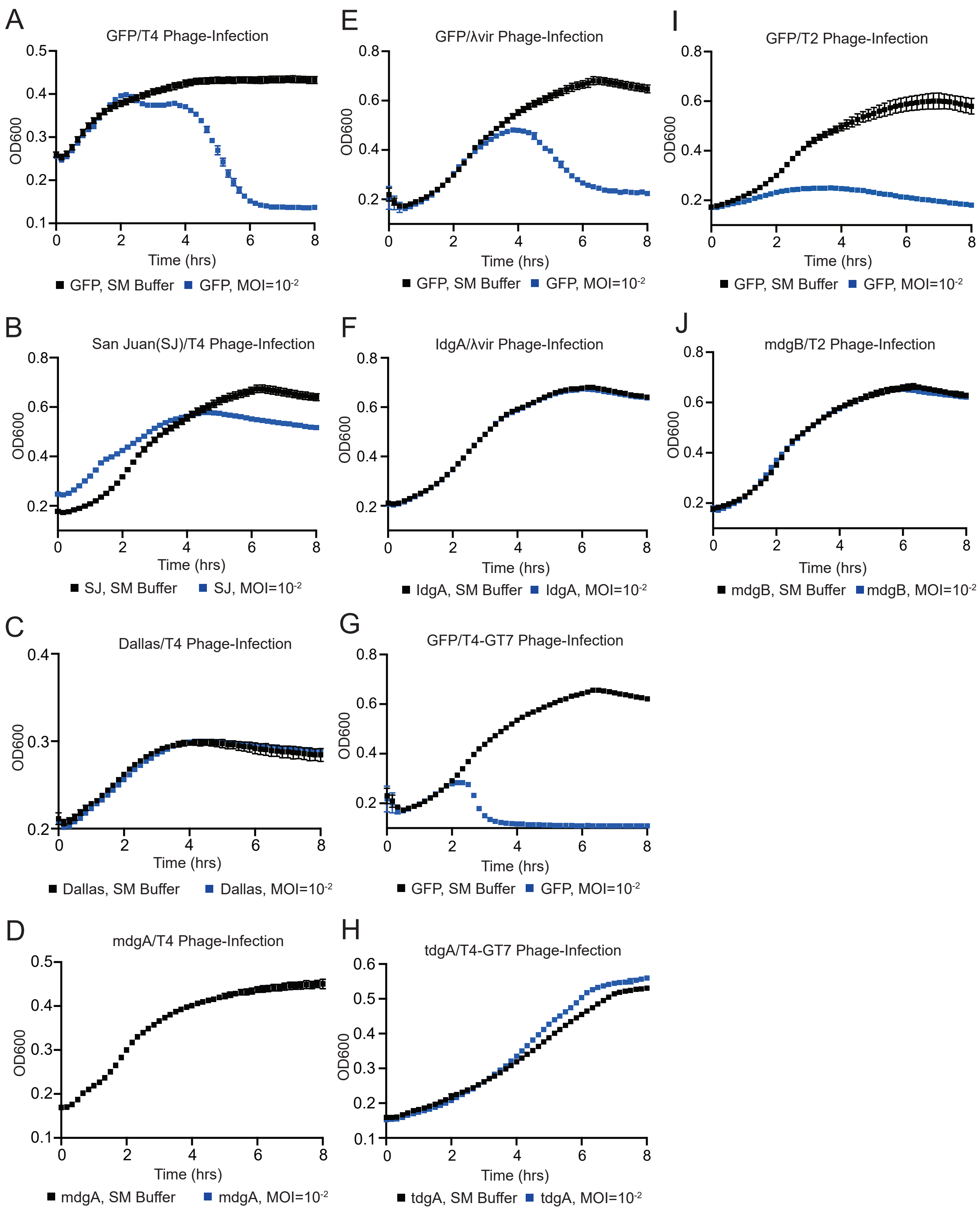


**Figure S6. Liquid phage infections with various defenses and controls.** (A-D) Bacterial growth curve with specified defenses without phage (SM Buffer) or with T4 phage added at MOI=$10^{-2}$. (A) GFP (B) San Juan (C) Dallas, and (D) *mdgA*. (E-F) Bacterial growth curve with specified defenses without phage (SM Buffer) or with $\lambda_{vir}$ phage added at MOI=$10^{-2}$. (E) GFP and (F) *ldgA*. (G-H) Bacterial growth curve with specified defenses without phage (SM Buffer) or with T4-GT7 phage added at MOI=$10^{-2}$. (G) GFP and (H) *tdgA*. (I-J) Bacterial growth curve with specified defenses without phage (SM Buffer) or with T2 phage added at MOI=$10^{-2}$. (I) T2 and (J) *mdgB*. The Y-axis label 'OD600' denotes the optical density readings at 600nm light (absorbance values).

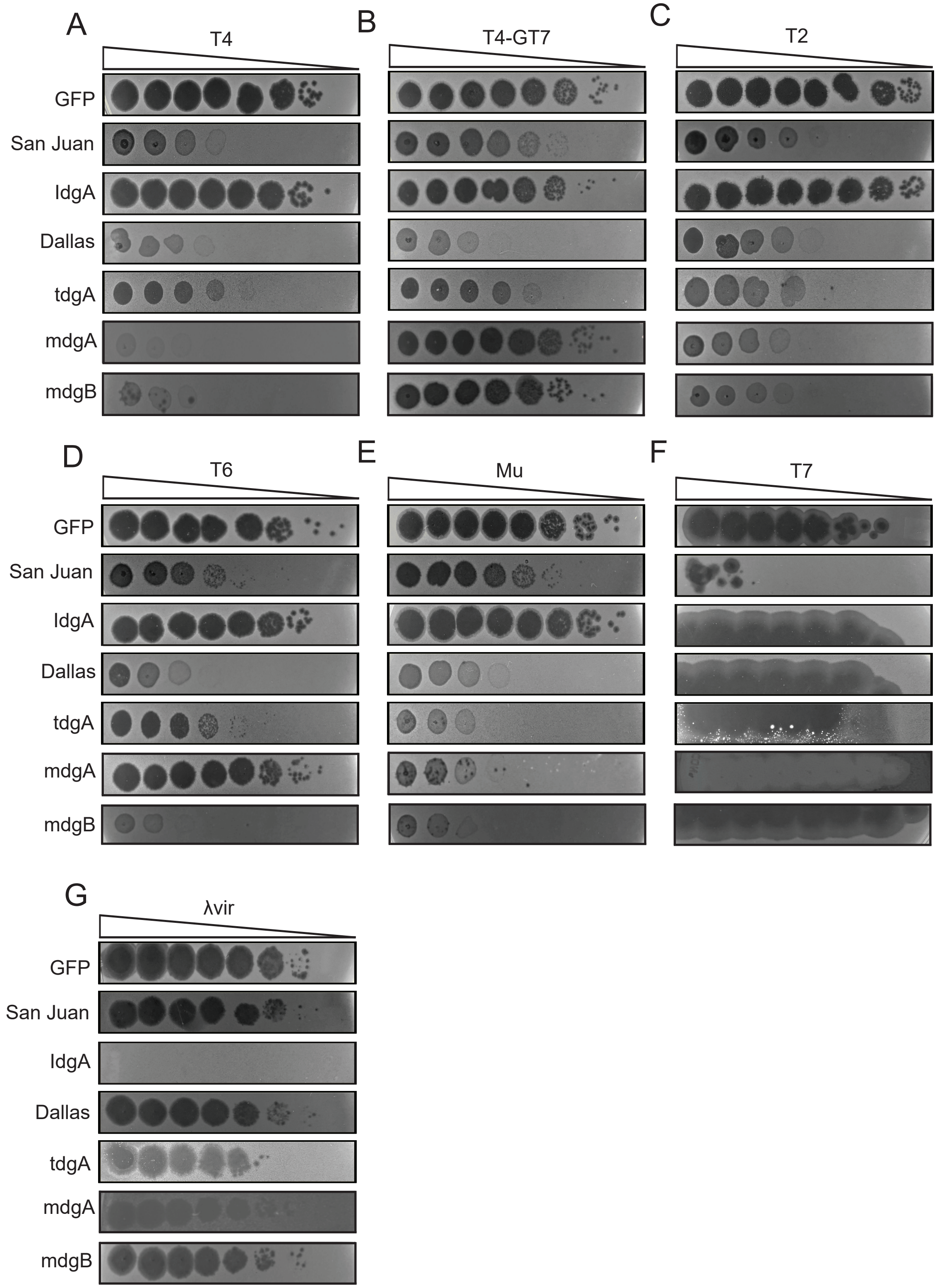


**Figure S7. Phage defense sufficiency experiments via plaque assays.** (A-G) GFP, San Juan, *IdgA*, Dallas, *tdgA*, *mdgA*, and *mdgB* subjected to solid-phage infection against (A) T4, (B) T4-GT7, (C) T2, (D) T6, (E) Mu, (F) T7 and (G) $\lambda_{vir}$. Experiments were performed in biological triplicate and representative plaque assays are shown.

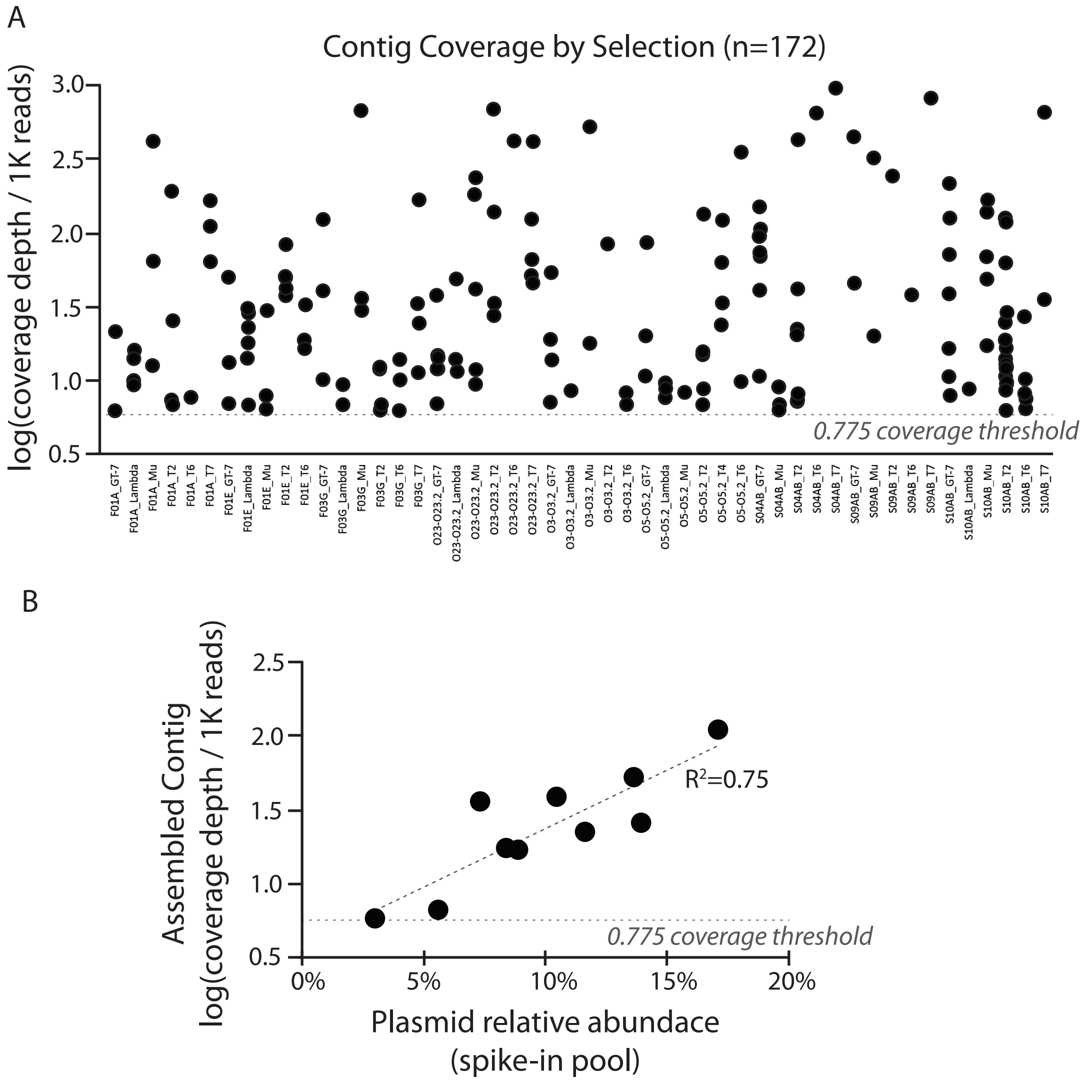


**Figure S8. Phage defense inserts are encoded by diverse metagenomes.** (A) Log-normalized coverage values for each contig assembled from pooled plasmid mixtures. Contigs from individually sequenced colonies are not included in this plot (all arise from T4 selections, see methods). The coverage threshold used as inclusion criteria is depicted as a dashed line. This coverage threshold corresponds to a plasmid at ~3% relative abundance in a control plasmid pool, per (B). (B) A defined mixture of confirmed phage-defense plasmids was sequenced alongside the selections depicted in (A). The relative abundance of each sequence cluster in the defined mixture correlated well with the log-normalized read coverage of the corresponding contig assemblies.
